# Arabidopsis REI-LIKE proteins activate ribosome biogenesis during cold acclimation

**DOI:** 10.1101/2020.02.18.954396

**Authors:** Bo Eng Cheong, Olga Beine-Golovchuk, Michal Gorka, William Wing Ho Ho, Federico Martinez-Seidel, Alexandre Augusto Pereira Firmino, Aleksandra Skirycz, Ute Roessner, Joachim Kopka

## Abstract

Arabidopsis REIL proteins are cytosolic ribosomal 60S-biogenesis factors. After shift to 10°C, *reil* mutants deplete and slowly replenish non-translating eukaryotic ribosome complexes of root tissue, while tightly controlling the balance of non-translating 40S- and 60S-subunits. *Reil* mutations compensate by hyper-accumulation of non-translating subunits at steady-state temperature; after cold-shift, a KCl-sensitive 80S sub-fraction remains depleted. We infer that Arabidopsis buffers fluctuating translation by pre-existing non-translating ribosomes before *de novo* synthesis meets temperature-induced demands. *Reil1 reil2* double mutants accumulate 43S-preinitiation and pre-60S-maturation complexes and have altered paralog composition of ribosomal proteins in non-translating complexes. With few exceptions, e.g. RPL3B and RPL24C, these changes are not under transcriptional control. Our study suggests requirement of *de novo* synthesis of eukaryotic ribosomes for long-term cold acclimation, feedback control of *NUC2* and *eIF3C2* transcription and links new proteins, AT1G03250, AT5G60530, to plant ribosome biogenesis. We propose that Arabidopsis requires biosynthesis of specialized ribosomes for cold acclimation.

**Highlight of this study:** REIL proteins affect paralog composition of eukaryotic ribosomes and suppress accumulation of 43S-preinitiation and pre-60S-maturation complexes, suggesting functions of ribosome heterogeneity and biogenesis in plant cold acclimation.

## Introduction

The *Arabidopsis thaliana* Col-0 (Arabidopsis) REI1-LIKE (REIL) proteins, REIL1 (At4g31420) and REIL2 (At2g24500) are homologs of the yeast Rei1 (YBR267W) and the paralog Reh1 (YLR387C) proteins. In yeast, both the Rei1 and Reh1 proteins function as ribosome biogenesis factors that participate either in parallel or sequentially in the late ribosome biogenesis step of cytoplasmic 60S ribosomal subunit maturation (Greber et al., 2012; Greber et al., 2016). Aside from this function, these proteins are required to maintain growth at suboptimal or cold temperatures. The *Δrei1* mutant is cold sensitive already at moderately suboptimal temperatures of yeast. The *Δrei1 Δreh1* double mutant is even more cold sensitive, while the yeast *Δreh1* mutation alone has no effect on growth in the cold (Iwase and Toh-e, 2004; Lebreton et al., 2006; Parnell and Bass, 2009). Heterologous expression of Arabidopsis REIL1, but not of REIL2 partly complements the cold sensitivity of the yeast *Δrei1* mutant (Schmidt et al., 2013). The Arabidopsis REIL paralogs differ in structure from their yeast homologs. REIL1 and REIL2 are four zinc finger proteins with C2H2 zinc finger domains arranged in two pairs, namely ZF1/ZF2 at the N-terminus and ZF3/ZF4 in the middle of the primary structure (Schmidt et al., 2013). In contrast, both yeast paralogs contain only three canonical zinc fingers (Iwase and Toh-e, 2004; Greber et al., 2012). This observation is in line with a preliminary phylogenic study (Schmidt et al., 2013), as both analyses indicate that the duplications of the yeast and Arabidopsis REIL paralogs are evolutionary independent (Schmidt et al., 2013).

Intriguingly, transfer-DNA (T-DNA) mutants of the Arabidopsis *REIL1* and *REIL2* genes, *reil1-1*, and allelic *reil2-1* or *reil2-2*, and the yeast *Δrei1* and *Δreh1* mutants have convergent growth responses at suboptimal temperature (Schmidt et al., 2013; Schmidt et al., 2014). Arabidopsis REIL paralogs are required specifically for plants to grow in the cold, e.g. at 4°C or at 10°C, but not for growth at optimized 20°C temperature (Schmidt et al., 2013; Beine-Golovchuk et al., 2018). In agreement with these findings, identical T-DNA insertion mutants of *reil2* were recently discovered in a screen for chilling sensitive Arabidopsis mutants (Yu et al. 2020), namely *reil2.1/ stch4-2* (GK_166C10) and *reil2.2/ stch4-1* (SALK_040068). The *reil1-1 reil2-1* double mutant has a more severe cold phenotype and all but stops development after germination at 10°C prior to the emergence of the first rosette leaf (Schmidt et al., 2013; Beine-Golovchuk et al., 2018). The double mutation is, however, non-lethal and a valid system for the investigation of REIL functions in plants. The double mutant maintains cellular integrity and acquires freezing tolerance after shifts from 20°C to 4°C or 10°C cold (Beine-Golovchuk et al., 2018). Expression of amino-terminal FLUORESCENT PROTEIN (FP)-REIL fusion proteins driven by the UBIQUITIN10 promoter rescue the mutant phenotypes (Beine-Golovchuk et al., 2018). The two allelic *reil2-1* and *reil2-2* mutants are both growth retarded and form small spoon-shaped leaves at 10°C (Schmidt et al., 2013; Yu et al., 2020). This phenomenon reverted after shift from cold to optimal 20°C. Except for slightly delayed germination, the *reil1-1* mutant, similar to yeast *Δreh1*, has no growth phenotype under the tested low temperature conditions (Schmidt et al., 2013; Beine-Golovchuk et al., 2018).

Previous functional analyses of Arabidopsis REIL proteins focused on vegetative photoautotrophic plant rosettes that were cultivated on soil and acclimated to 10°C temperature. Conserved functions of the Arabidopsis REIL proteins and new functions that evolved on the path towards multicellular embryophyte plants became apparent (Beine-Golovchuk et al., 2018). As was expected in analogy to yeast, FP-REIL1 and FP-REIL2 fusion proteins localize to the cytosol (Beine-Golovchuk et al., 2018; Yu et al., 2020) and the REIL1 protein appears to interact with ribosome complexes containing the 60S large ribosome subunit (LSU). Based on immunoprecipitation-mass spectrometry, REIL2 associates with the eukaryotic ribosome (Yu et al., 2020). Our current study locates REIL2 to the non-translating 60S fraction of Col-0 wild type.

In rosette leaves, REIL proteins accelerate 10°C cold induced accumulation of cytosolic ribosome subunits and of cytosolic ribosomal RNA (rRNA). After cold shift, they enhance gene expression of structural proteins of cytosolic ribosomes, ribosome biogenesis factors, and cytosolic translation initiation or elongation factors (Beine-Golovchuk et al., 2018). The acclimation responses occur within the first week after cold shift, while Arabidopsis wild type plants pass through a lag-phase before resuming growth. All of these processes lag behind in the *reil1-1 reil2*-*1* mutant that does not resume growth in the cold (Schmidt et al., 2013). Our current study confirms these observations in roots and provides mechanistic insights far beyond. Besides an influence on cold-induced plant ribosome remodeling and on the accumulation of cytosolic ribosome subunits after cold shift, Arabidopsis REIL proteins are apparently involved in plant-specific processes. For example, the *reil1-1 reil2-1* mutant does not activate FLOWERING LOCUS T gene expression in mature leaves after the cold shift and exhibits plant cold-acclimation responses at the 20°C non-acclimated state, including premature activation of the C-REPEAT/DRE BINDING FACTOR1/DEHYDRATION-RESPONSIVE ELEMENT BINDING (CBF/ DREB) regulon (Beine-Golovchuk et al., 2018). Analyses of the single paralog *reil2/ stch4* mutants confirmed the interaction of *reil2* function with the CBF/ DREB regulon, e.g. (Cook et al., 2004; Maruyama et al., 2004; Van Buskirk and Thomashow 2006), with REIL2 deficiency delaying CBF/ DREB regulon activation and reducing CBF/ DREB protein accumulation in the cold. Ectopic overexpression of REIL2 under control of the 35S promoter conveyed enhanced chilling and freezing tolerance under mixotrophic *in vitro* conditions, i.e. in the presence half-strength Murashige and Skoog (MS) medium (Yu et al., 2020).

To reveal organ-independent and likely direct or primary functions of Arabidopsis REIL proteins from more indirectly associated responses to REIL deficiency, we analyzed the root system of Arabidopsis Col-0 wild type and of *reil* mutant plants under 20°C to 10°C cold shift conditions and compared these results to previous analyses of mutant and wild type Arabidopsis rosette leaves. To obtain sufficient amounts of root material for multi-level systems analyses, we changed our experimental conditions from soil-grown plants (Beine-Golovchuk et al., 2018) to a mixotrophic hydroponic system that meets these demands. In this cultivation system, *reil* mutants retain reduced growth, but attenuate the previously reported strong growth and developmental phenotypes. For our current study, we did not adjust the growth conditions to reveal stronger phenotypes. By analyzing *reil* mutants under conditions that cause small, but still noticeable phenotypes, we hope to reveal the mutant effects that otherwise may be obscured by pleiotropic and secondary responses. For the purposes of: (1) identifying and understanding the mechanism of ribosome biogenesis in which Arabidopsis REIL proteins are involved and; (2) discovering alternative roles of REIL proteins, we analyzed ribosomal complexes and applied transcriptome and proteome profiling methods. In addition to the previously reported double mutant *reil1-1 reil2*-*1* (Schmidt et al., 2013; Schmidt et al., 2014; Beine-Golovchuk et al., 2018), we introduce a second double mutant, *reil1-1 reil2-2*, to our studies and analyzed abundances of ribosomal complexes from the single-paralog mutant lines, *reil1-1*, *reil2-1* and *reil2-2*, from which the double mutants originate.

## Materials and methods

### Plant material

The single-paralog mutants, *reil1-1*, *reil2-1*, and *reil2-2* lines, with T-DNA insertions in exon 2 at base pairs 475, 733, and 731, respectively, were selected to investigate the role of REIL proteins for the plant growth in suboptimal temperature condition. All T-DNA insertions were characterized previously by sequencing of the left border of insertion sites (Schmidt et al., 2013). The double homozygous *reil1-1 reil2-1* (*DKO1*) mutant (Schmidt et al., 2013) was created by crossing the T-DNA insertion mutant SALK_090487 (*reil1-1*) that was obtained through the Nottingham Arabidopsis Stock Centre (Scholl et al., 2000) and GK_166C10 (*reil2-1*) of the GABI-Kat program (Rosso et al., 2003). The second allelic double homozygous *reil1-1 reil2-2* (*DKO2*) mutant was created in this study by crossing the T-DNA insertion mutant SALK_090487 (*reil1-1*) and *reil2-2* (SALK_040068). Homozygosity of the insertion sites was verified by PCR amplification of genomic DNA with the previously described T-DNA- and *reil1*- and *reil2*-specific oligonucleotides (Schmidt et al., 2013). All system-profiling analyses were performed with the two double mutants and controls of the ecotype *Arabidopsis thaliana* Col-0 (wild type), the common genetic background of all mutants in this study. The three single-paralog *reil* mutants, *reil1-1*, *reil2-1* and *reil2-2*, were subjected to comparative analysis of ribosome complexes.

### Photographic documentation

The Arabidopsis shoot and root systems were digitally photographed at 300 dpi horizontal and vertical resolution using a Nikon D5100 camera with up to 4,928 x 3,264 pixels or a NIKON D850 camera with up to 8,256 x 5,504 pixels. Photographs were cropped to size and resolution reduced to meet size limitations of composite figures using Adobe Photoshop CS5 Extended (Version 12.0.4 x64). The scale bars were manually added using external documentation of scale and magnification.

### Hydroponic system for growth, controlled temperature shift and sampling of Arabidopsis root materials

Generation of seed batches, storage, sterilization, and *in vitro* pre-cultivation were as described previously (Schmidt et al., 2013). The hydroponic growth system for Arabidopsis plants used circular white glass containers with glass lids. The containers had a slightly tapered bottom with 10.0 cm top diameter and 7.5 cm bottom diameter (**Supplemental Fig. S1**). The containers had a total volume of ∼ 580 mL. Each glass contained 250 mL of liquid MS media with 2% sucrose (w/v) adjusted to pH 5.7 (Murashige and Skoog, 1962). Sucrose addition reduced developmental and growth-differences between mutants and Col-0 wild type. The liquid was first poured into the jar, then a sterilized stainless steel wire mesh insert of 9 cm diameter, with mesh size 1.40 mm and wire thickness 0.25 mm was adjusted level with the top of the liquid (**Supplementary Fig. S1**). Four small ∼ 1 cm^3^-blocks of solid MS media that contained 0.8 % (w/v) agar were placed on top of the mesh before applying single sterilized Arabidopsis seeds to each block. Four plants were routinely cultivated per container. The containers were closed tightly, but not airtight. Glass lids were fixed to the container by stainless steel clamps before transfer to a controlled-environment chamber for growth. The seeds were allowed to germinate and grow under sterile conditions in a long day photoperiod with a 16h/8h - day/night cycle. External temperature settings of the chamber were controlled. External light intensity was 100 - 150 µmol photons m^-2^ s^-1^. External relative humidity was 60% - 70%. The standard temperature regime was 20°C during the day and 18°C during night (**Supplementary Fig. S1**).

### Temperature shift experiments and root harvests

Plants were pre-cultivated at 20°C standard conditions to developmental stage ∼1.10 (Boyes et al. 2001). At this stage, the plants were transferred carefully within their hydroponic containers from standard conditions to a second controlled-environment chamber with 10°C temperature during the day and 8°C during the night, but otherwise equal external light cycle, light intensity, and relative humidity settings (**Supplemental Fig. S2**). Non-cold shifted roots were harvested immediately before temperature shift and are in the following denoted non-acclimated or 0 day samples. Subsequently, samples were harvested after one day, three days, one week, and three weeks after cold shift (**Supplemental Fig. S3**). Each harvested biological replicate was the pool of total root material from four plants of a single container. Three biological replicates, n = 3, were harvested for microarray based transcriptome analyses. Five replicates per experimental condition, ∼20 individual root systems, were pooled into single samples for the analysis of ribosome complexes using sucrose density gradients.

### Sucrose density gradient analysis of ribosome preparations from Arabidopsis root tissue

Ribosome fractions from Arabidopsis Col-0 wild type and *reil* mutants were prepared from pools of hydroponically grown root systems as described previously (Beine-Golovchuk et al., 2018). Due to the complex procedure, single pools of non-acclimated roots and roots at 1 day, 3 days, 7 days, and 21 days after cold-shift were analyzed per genotype for qualitative analyses. Briefly, per sample 100 - 102 mg fresh weight (FW) of frozen root tissue was homogenized 30 min on ice with 0.5 mL polysome extraction buffer (PEB) in horizontal tubes on an orbital shaker set to 400 rotations per minute. The final PEB buffer composition was 200 mM Tris-HCl adjusted to pH 9, 200 mM KCl, 25 mM EGTA, 36 mM MgCl_2_, 5 mM DTT, 50 µg mL^-1^ cycloheximide (CHX), 50 µg mL^-1^ chloramphenicol (CHL), 1.0 mg mL^-1^ heparin, 1 % (v/v) Triton X-100, 1 % (v/v) Tween 20, 1 % (w/v) Brij-35, 1 % (v/v) IGEPAL CA-630, 1 % (v/v) polyoxyethylene, 1 % (w/v) deoxycholic acid, 1 mM phenylmethylsulfonylfluoride (PMSF), and the equivalent of 0.002 tablets mL^-1^ cOmpleteTM protease inhibitor mixture. All chemicals were obtained from Sigma-Aldrich (Taufkirchen, Germany). The following procedures were carried out at 4°C. After initial centrifugation at 14,000 g for 10 min, the extract with remaining debris was placed on top a QIAshredder spin column of the RNeasy Plant Mini Kit (Qiagen, Hilden, Germany) and centrifuged again at 14,000 g for 1 min to remove all cell debris. Volumes of 0.5 mL of extract were loaded onto 9 mL of a 15% (w/v) to 60% (w/v) sucrose gradient that was prepared in PEB without detergents, heparin and protease inhibitors. The loaded density gradients were centrifuged at 33,000 g for 14.5 h at 4°C using a six-position SW41 Ti rotor (Beckman Coulter, Kerfeld, Germany). The positions of ribosome complexes varied slightly with each centrifugation run that comprised a set of six samples. In each centrifugation run, a non-sample control gradient was loaded with 0.5 mL PEB and used for photometric base line subtraction. The five remaining positions were used for root samples and included an Arabidopsis Col-0 sample for the estimation of relative abundances of ribosome complexes. Five centrifugation runs contained samples that were harvested at the same experimental time point. In the sixth centrifugation run, a randomly selected mutant of each time point was analyzed. The sucrose density gradients were separated into ∼40 fractions of approximately 250 µL per fraction using a programmable density gradient fractionation system (Teledyne Isco Inc., NE, USA). An absorbance profile was recorded continuously at 254 nm wavelength using a 15% (w/v) sucrose solution in PEB without detergents, heparin and protease inhibitors, prior to each gradient recording to calibrate the absorbance minimum (0% absorbance_254nm_). The absorbance profiles were exported from the acquisition system and background subtracted by the profile of blank sample of each centrifugation run and identically scaled. Abundances of ribosome fractions were measured as background subtracted peak areas (**Supplemental Table S1**). Peak areas and the total sum of all peak areas including the 40S, 60S, 80S and polysome fractions, were determined by Chromas Lite 2.1 software (http://chromas-lite.software.informer.com/2.1/).

### KCl-sensitivity test of ribosome complexes

Exemplary *in vitro* KCl-sensitivity of translating ribosome complexes was performed with wild type root tissue compared the *reil1-1 reil2-1* double mutant at 21 days after cold shift as a compromise between immediate and prolonged cold exposure. A pooled root sample of each simple type was split into two technical replicates of 100 mg FW each. Each of the replicates was extracted with 0.5 mL PEB that either contained regular 200 mM KCl or elevated 400 mM KCl. After 30 min incubation on ice all samples were fractionated by 15% - 60% sucrose density gradients. All samples were centrifuged in a single 2 h run with sedimentation at 50,000 g and 4°C using small 13 x 51 mm ultracentrifugation tubes in SW55 Ti rotor (Beckman Coulter, Krefeld, Germany). These centrifugation settings were optimized for polysome and monosome separations. Abundance analysis of the separated ribosome complexes was as described above omitting fraction collection.

### Transcriptome analysis

Twenty seven root samples, 3 experimental conditions x 3 genotypes x 3 biological replicates, of non-acclimated and 1 day or 7 days cold acclimating Col-0, *reil1-1 reil2-1*, *reil1-1 reil2-2* were analyzed. Liquid nitrogen frozen samples of 25.1 - 26.8 mg fresh weight were ground to fine powder. Total RNA was extracted from frozen powder using the RNeasy Plant Mini Kit and RNase-free DNase (Qiagen, Limburg, Netherlands) according to the manufacturer’s instructions as was described previously (Beine-Golovchuk et al., 2018) with down-scaling modifications. Briefly, 110 µL RNA extraction buffer was used and 30 µL of RNase-free water for RNA elution. Total RNA was quantified by NanoDrop™ spectrophotometer (ThermoFisher Scientific, Germany) using absorbance wavelength 260 nm. RNA quality was evaluated by formaldehyde agarose gel electrophoresis and OD_260/280_ and OD_260/230_ ratios. The RNA integrity number (RIN) of RNA samples was assessed using an Agilent 2100 Bioanalyzer (Agilent Technologies, Germany). RNA samples of 10-15 µL per sample with RIN > 8 were processed by ATLAS Biolabs GmbH (Berlin, Germany) using Agilent Feature Extraction software (v10.7). The obtained gene expression data sets were quantile normalized using the normalize.quantile routine of the preprocessCore R statistical computing and graphics package (https://www.r-project.org/). The obtained 4X44K Agilent microarray data comprise variable numbers of redundant gene probes. The expression data sets are available from the Gene Expression Omnibus (https://www.ncbi.nlm.nih.gov/geo/) through accession GSE144916. Gene expression information of redundant probes was averaged to reduce bias for functional enrichment analyses. Gene expression information was normalized per gene model to the average expression value from non-acclimated Col-0. Alternatively, the differential gene expression of mutants over Col-0 was calculated at each time point. Averaged log_2_-transformed ratios of differential gene expression, standard errors and statistical test results are reported in the supplement (**Supplemental Table S2**).

### Functional enrichment analyses of gene expression data

Functional enrichment analysis of differential gene expression was carried out by parametric analysis of gene set enrichment (Du et al., 2010; Tian et al., 2017) using the agriGO v2.0 web-service at http://systemsbiology.cau.edu.cn/agriGOv2/. The parametric analysis of gene set enrichment processes lists of genes with log_2_-transformed numerical differential gene expression values from one or more experimental conditions and provides information on the enrichment of expression increases or decreases across gene sets of at least 10 gene entries that are defined by 2,145 GO gene ontology (GO) terms. Enrichment of mean log_2_-fold changes (log_2_-FC) were evaluated by z-scores, and by false discovery rate (FDR)-adjusted *P*-values (Benjamini and Hochberg, 1995) applying a *P* < 0 .05 threshold (**Supplemental Table S3**).

### Proteome Analysis of ribosome preparations from Arabidopsis root tissue

We performed two complex experiments of ribosome fraction preparation and shotgun ribo-proteome analysis that resulted in data sets DS1 and DS2, respectively. Each data set contained a pooled Arabidopsis Col-0 root sample and a pooled double mutant root sample either of *reil1-1 reil2-2* (DS1) or of *reil1-1 reil2-1* (DS2). Samples prior to temperature shift and samples prepared at 7 days after 10°C cold shift were analyzed.

Sucrose density fractionation and subsequent proteomic analysis were performed with approximately equal amounts of 100 mg FW. Tryptic peptides were prepared from up to five non-translating ribosome fractions and a low-oligomer polysome fraction by filter aided sample preparation (Erde et al., 2014; Swart et al., 2018). Sucrose density gradient fractions were washed repeatedly with 500 µL of 0.04 M Tris-HCL buffer (pH 8.4) with 0.2 M KCl and 0.1 M MgCl_2_ using regenerated cellulose membranes, Amicon Ultra-0.5 centrifugal filter units, with a 3 kDa molecular size cutoff (Merck, Kenilworth, New Jersey, USA). Removal of sucrose was considered complete, when the 500µL volume decreased to below 100 µl within 10 minutes of centrifugation at 5000-7000 rpm and 4°C.

Shotgun proteome analysis of recovered peptides was performed by liquid chromatography - tandem mass spectrometry (LC-MS/MS) with an ACQUITY UPLC M-Class system (Waters Corporation, Milford, MA, USA) hyphenated to a Q-Exactive HF high-resolution mass spectrometer (Thermo Fisher Scientific, Waltham, MA, USA). Samples were separated by reverse-phase nano-liquid chromatography using a 125/ 132 min (DS1/ DS2) 3% to 85% (v:v) acetonitrile (ACN) gradient. Mass spectrometric data acquisition was performed by data dependent top-N tandem mass spectrometry (dd-MS2). This acquisition mode fragmented the top 10/ 15 (DS1/ DS2) most intense ions per full scan. Full scans were acquired at a resolution of 120,000/ 60,000 (DS1/ DS2) with automatic gain control (AGC) target set to 3e6/ 1e6 (DS1/ DS2), maximum injection time 100/ 75 ms (DS1/ DS2), and scan range 300 to 1600 m/z in profile mode. Each dd-MS2 scan was recorded in profile mode at a resolution of 15,000/ 30,000 (DS1/ DS2) with AGC target set to 1e5, maximum injection time 150 ms, isolation window of 1.2/ 1.4 m/z (DS1/ DS2), normalized collision energy 27 eV and dynamic exclusion of 30 sec. The LC-MS/MS files were analysed by MaxQuant software (Version 1.6.0.16), http://www.coxdocs.org/doku.php?id=maxquant:common:download_and_installation. A*rabidopsis thaliana* FASTA files of the reviewed Swiss-Prot compendium in the UniProt database (The UniProt Consortium, 2017) served peptide annotation. All peptides, including unique peptides and non-specific peptides of highly similar or identical paralogs of structural cytosolic ribosome proteins were analyzed by label-free quantification, LFQ (Zhang et al., 2012).

The proteomics data set, identifier PXD016292, and detailed experimental settings are available via the ProteomeXchange Consortium at the PRIDE partner repository, https://www.ebi.ac.uk/pride/archive/login, (Perez-Riverol et al., 2019). Proteins were annotated according to majority peptide identification. Specificity of detected peptides for cytosolic RP paralogs and ribosome-associated proteins (RAPs) was analyzed including potential non-specificity by minority peptide calls. Supplemental Table S4 reports proteomic fraction characterization and the subset of samples from Pride data set PXD016292 used in this study. We identified the 40S and 60S RPs, and the organelle 30S and 50S RPs according to previous publications (Barakat et al., 2001; Hummel et al. 2015; Sormani et al., 2011; Waltz et al., 2019; Rugen et al., 2019). Due to the differences of cytosolic ribosome subunits in each fraction, and to account for the variation between sample types, we normalized protein LFQ-abundances in fractions that contained 60S RPs to the sum of LFQ-abundances of all detected 60S proteins in each of the respective fractions. This procedure allowed analysis of compositional changes of 60S ribosome complexes, but may create false positives of components due to co-purified organelle RPs or other proteins. For this reason, we only considered changes relevant, if the non-normalized LFQ abundance had a maximum in the 60S or 60S/80S fractions and not in the 30S/40S or 50S fraction. We excluded all 40S RPs from the compositional analysis of the non-translating 60S fractions. To assess compositional changes of non-translating 40S complexes we normalized to the sum of all 40S RPs in each fraction. We excluded all 60S RPs and only considered proteins that we more abundant in the 30S/40S fraction relative to the 60S fraction. Due to the overall experimental design, we did not apply statistical analyses for feature selection. Instead, we selected for changes that were shared between the *reil1 reil2* double mutants at 10°C. In a more stringent analysis, we selected the respectively shared changes at both temperatures 10°C and 20°C.

### Association analysis of changes in transcript and relative protein abundance

The differential transcript levels of the *reil1 reil2* mutants relative to *Arabidopsis thaliana* Col-0 wild type were compared to changes of relative protein abundance in the non-translating 40S and 60S plus 60S/80S fractions of the proteome analysis (**Supplemental Figure S9**). For this purpose, we correlated the differential accumulation of cytosolic RPs and RAPs in non-translating ribosome fractions reported in Supplemental Table S6 to differential gene expression (**Supplemental Table S2**, columns X to AC). We merged transcript and protein information according to the reported gene model information of the transcriptome and proteome analyses (Pride data set PXD016292). Note that the transcript information was in part splice variant specific. The proteome analyses in many cases did not distinguish splice variants and contained non-specific information due to peptides of proteins from the highly similar or in part identical ribosome protein families (**Supplemental Table S6**).

### Statistical analyses and data visualizations

Quantile-normalized log_2_-transformed ratios of DEGs (differentially expressed genes) or log_2_-transformed of differential protein abundances were analyzed, correlated and visualized by Microsoft Excel software of the Office Professional Plus 2010 package (Microsoft) and the Multiple Experiment Viewer version 4.9.0 (http://www.mybiosoftware.com/mev-4-62-multiple-experiment-viewer.html). Significance thresholds (*P*) of statistical analyses, e.g. one- or two-way analyses of variance (ANOVA), and two-group comparisons by heteroscedastic Student’s t test, were routinely *P* < 0.05 if not mentioned otherwise. Details of the data analysis and visualization methods are reported in the legends of figures and tables.

## Results

### Hydroponic growth of *reil* mutant plants compared to the Arabidopsis Col-0 wild type

The established hydroponic system enabled sterile photomixotrophic cultivation of up to four Arabidopsis Col-0 wild type or mutant plants within a single container with 2% (w/v) sucrose in liquid growth medium (Murashige and Skoog, 1962). Attempts of photoautotrophic cultivation without sucrose caused slow plant growth and generated dwarfed and early flowering *reil1 reil2 double* mutants (DKOs) with minimal root systems. In the presence of 2% (w/v) sucrose, the Arabidopsis wild type reached the 10-leaf stage, i.e. Arabidopsis stage 1.10 (Boyes et al., 2001), after approximately 4 weeks under standard long-day photoperiod conditions with external temperatures set to 20°C during the day and 18°C during night (**Supplementary Fig. S1**). Morphology of Arabidopsis rosettes differed from soil-grown plants. The leaf petioles were elongated and leaf laminae smaller than on soil, e.g. (Schmidt et al., 2013; Beine-Golovchuk et al., 2018). Leaves of individual stage 1.10 plants overlapped minimally. From this stage onward, the single root systems entangled with each other and in part with the mesh. Only rapid harvests of the complete root systems from four plants of a single container were feasible (**Supplementary Fig. S2**). For this reason, all biological replicates were pooled root samples of at least four plants from single growth containers or of more plants from multiple containers.

Root and shoot growth of the Arabidopsis wild type continued after cold shift at stage 1.10 plants to 10°C during the day and 8°C during night under otherwise constant conditions. Plants phase-shifted to the generative phase and produced small inflorescences at 7 to 21 days after exposure to cold (**Supplementary Fig. S2**). Photomixotrophic cultivation did not fully abolish differential growth of mutants and Col-0. Growth and development of the *reil1-1* mutant was, however, similar to wild type. Shoots of the mutants, *reil2-1, reil2-2,* and the double mutants, *reil1-1 reil2-1 (DKO1)* and *reil1-1 reil2-2 (DKO2)* remained slightly smaller than wild type after cold shift, but all mutants continued to grow and developed inflorescences (**Supplementary Fig. S2**). Shift to low temperature induced spoon-shaped leaf morphology of *reil2* mutants, but this phenotype was strongly attenuated compared with soil-grown *reil2* mutants (Schmidt et al. 2013). Similarly, the double mutants were not fully growth arrested after 10°C cold shift in our hydroponic growth system and continued to grow in contrast to soil cultivation (Beine-Golovchuk et al., 2018).

A preliminary study of mutant root systems from single plants of our cultivation system (**Supplementary Fig. S3**) showed delayed root growth and a shortened acropetal part of the primary root of the double mutants compared with the wild type. These observations became apparent 7 to 21 days after cold shift. In addition, at 21 days after cold shift, the double mutants and *reil1-1* appeared to be less branched, indicative of an altered root branching pattern compared to the wild type (**Supplementary Fig. S3**). Attempts to further characterize altered root growth and branching pattern without wounding failed in our current cultivation system because it was inadequate to quantify altered root architectures of single plants at stage 1.10 and later. For this reason, we did not perform morphometric analyses of mutant root systems in this study. In the following, we performed standardized temperature shift experiments that were similar in experimental design to previous experiments with soil-grown plants (Schmidt et al., 2013; Beine-Golovchuk et al., 2018). We analyzed pools of whole root systems and started experiments at developmental stage 1.10 with samples taken in the non-acclimated state followed by sampling at 1, 3, 7, and 21 days after shift to 10°C cold.

### Analyses of ribosome complex abundances from roots confirms association of REIL deficiency with delayed 60S LSU accumulation after cold shift

Previous analyses of cytosolic ribosome complexes from Arabidopsis showed activation of cytosolic rRNA expression and concerted accumulation of transcripts coding for cytosolic ribosomal proteins (RPs) after a 10°C cold shift from optimized temperatures (Beine-Golovchuk et al., 2018). The *reil1-1 reil2-1* mutant delayed these responses and the accumulation of the 60S LSU fraction compared to wild type. In these studies, we analyzed leaf material of soil-cultivated rosette plants. The presence of chloroplast ribosomes interfered in parts with the analysis of cytosolic ribosome complexes. We succeeded to separate a 60S LSU fraction from the chloroplast 50S LSU and chloroplast 70S from 80S monosomes, but the chloroplast 30S and cytosolic 40S small subunits (SSUs) remained non-resolved and the polysome fraction was mixed. For this reason, we did not investigate previously, an indicated concerted regulation of cytosolic 60S LSU with the abundances of other cytosolic ribosomes complexes, such as the 40S SSUs.

To overcome this limitation and to extend and validate our analyses of the delayed accumulation of the 60S LSU in REIL deficient mutants, we focused on root material that became available in sufficient amounts, ∼ 100 mg fresh weight per analysis, by hydroponic cultivation. As we expected, plastid ribosome complexes, 30S, 50S, and 70S, were negligible in this material and much less abundant than cytosolic ribosomes. We obtained improved separation and purity of 40S SSU, 60S LSU, 80S ribosome fractions and analyzed the abundance of these fractions in wild type compared to the double mutants, *reil1-1 reil2-1* and *reil1-1 reil2-2*, and the respective single-paralog mutants, *reil1-1*, *reil2-1*, and *reil2-2* (**Fig. 1**). Based on the equal amounts of root fresh weight Arabidopsis wild type accumulated cytosolic ribosome complexes, especially the 80S fraction, at 1, 3, 7, and 21 days after shift to 10°C. Profiles of ribosome complexes from both, the *reil1-1 reil2-1* and the *reil1-1 reil2-2* mutant, were distinct from wild type. The abundance of the 60S LSU fraction of both double mutants approximately equaled the 80S fraction already in the non-acclimated state. This difference to wild type persisted throughout cold acclimation (**Fig. 1**). In agreement with previous observations from leaf material, 60S LSU accumulation after cold shift lagged behind wild type in both *reil* double mutants.

**Figure 1.**
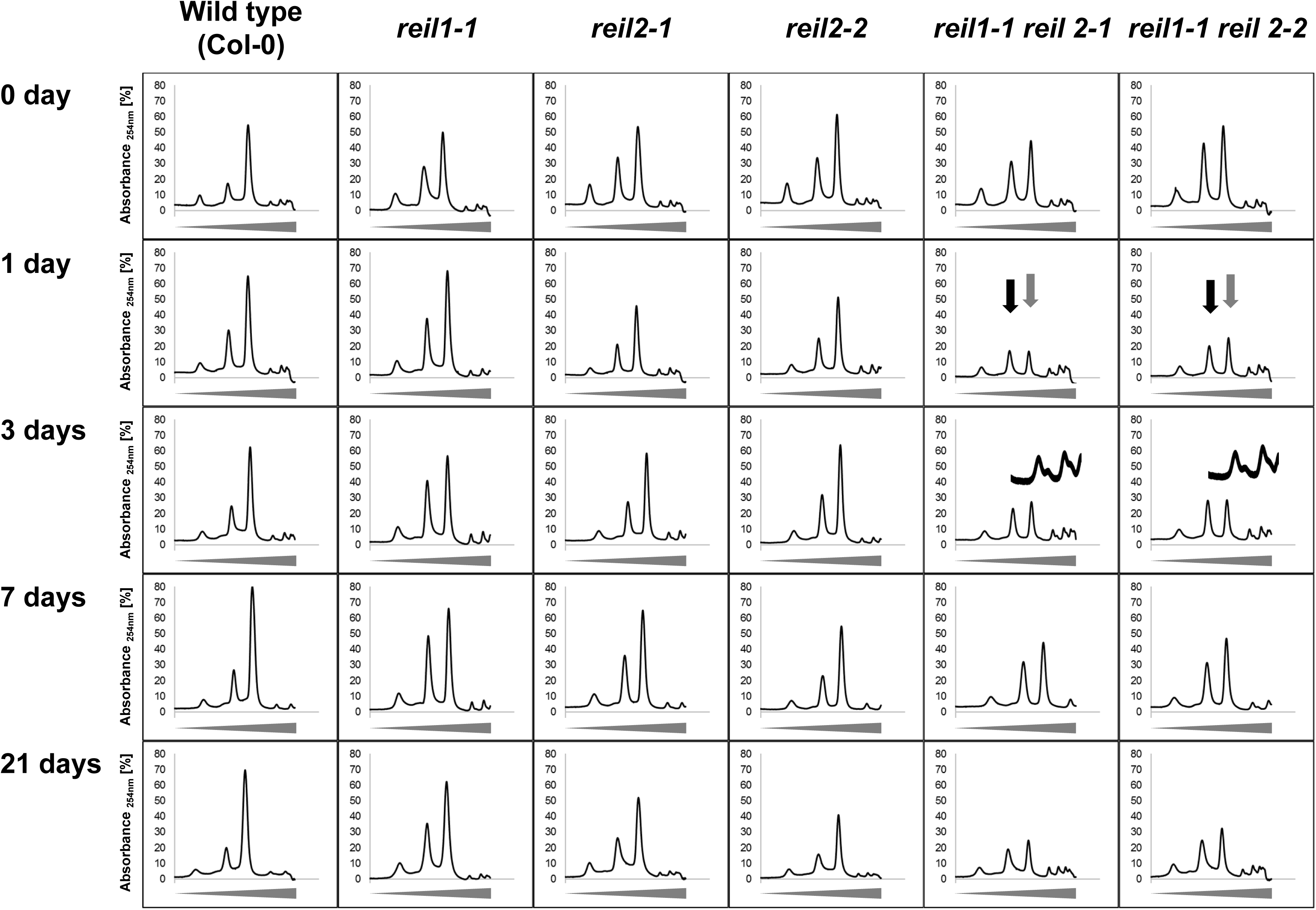
Sucrose density gradient analyses of ribosome complexes from equal fresh weight amounts of hydroponically grown total root material before (0 day) and at 1, 3, 7, or 21 days after shift from 20°C (day)/ 18°C (night) to 10°C (day) and 8°C (night). The *Arabidopsis thaliana* wild type (Col-0) was compared to the single-paralog mutants, *reil1*-*1*, *reil2*-*1*, *reil2*-*2*, and to the double mutants, *reil1*-*1 reil2*-*1* and *reil1*-*1 reil2*-*2*. Absorbance at wavelength 254 nm (y-axis) was recorded continuously during fractionation of 15 – 60 % sucrose density gradients (x-axis, tapered bars indicate the gradient) and background corrected by a non-sample control. This is a composite figure of multiple centrifugation runs that each contained a non-sample control. Gradients varied slightly with each centrifugation run. Note the altered abundance pattern of ribosome complexes from the *reil1*-*1 reil2*-*1* and *reil1*-*1 reil2*-*2* double mutants in non-acclimated and acclimating states and the delay relative to Col-0 of 60S LSU (black arrows) and 80S monosome (grey arrows) accumulation after cold shift. Inserts into the day 3 analyses magnify the low-oligomer polysome region and demonstrate accumulation of half-mer polysomes in the double mutants early after cold shift.

### Changes of ribosome complex abundances reveal inter-complex correlations and compensation responses to REIL deficiency

This observation and associated effects of other ribosome fractions became apparent by analyzing log_2_-transformed ratios of ribosome complex abundances of mutants compared to wild type at each time point of our temperature shift experiment (**Fig. 2, Supplemental Table S1**). In the non-acclimated state, all *reil* mutants increased the 60S LSU and the 40S SSU fractions ∼two-fold relative to wild type (**Fig. 2A-B**), but 80S monosomes had wild type levels (**Fig. 2C**). These observations in the non-acclimated state coincided with a slight increase of sum of all detected ribosome complexes (**Fig. 2D**). At one day after cold shift, all *reil* mutants except *reil1-1* reduced the 60S LSU and the 40S SSU fractions to approximate wild type levels (**Fig. 2A-B**). Reductions of these fractions were only marginal in the *reil1-1* mutant. All mutants subsequently returned to at least wild type levels and in most cases approximated the initial over-accumulation of 60S LSUs and 40S SSUs. The *reil1-1 reil2-1* and the *reil1-1 reil2-2* double mutants reduced the abundance of the 80S fraction up to four-fold relative to wild type upon cold shift (**Fig. 2C**). This phenomenon occurred at 1 day after cold shift and continued during prolonged cold exposure. In the *reil2* single-paralog mutants, the relative abundances of the 80S fraction were similar, but less affected. The sum of all ribosome complexes in the double mutant changed only marginally, but was reduced at day 1 and 21 after cold shift in the double mutants and in part in the single mutants (**Fig. 2D**). The changes of 60S LSU and the 40S SSU abundances correlated with a Pearsońs correlation coefficient of r = 0.945 across all measurements of our experiment (**Fig. 3A**). The relative abundances of the 60S LSU and 80S monosome fractions, however, correlated less stringently, r = 0.607 (**Fig. 3B**).

**Figure 2.**
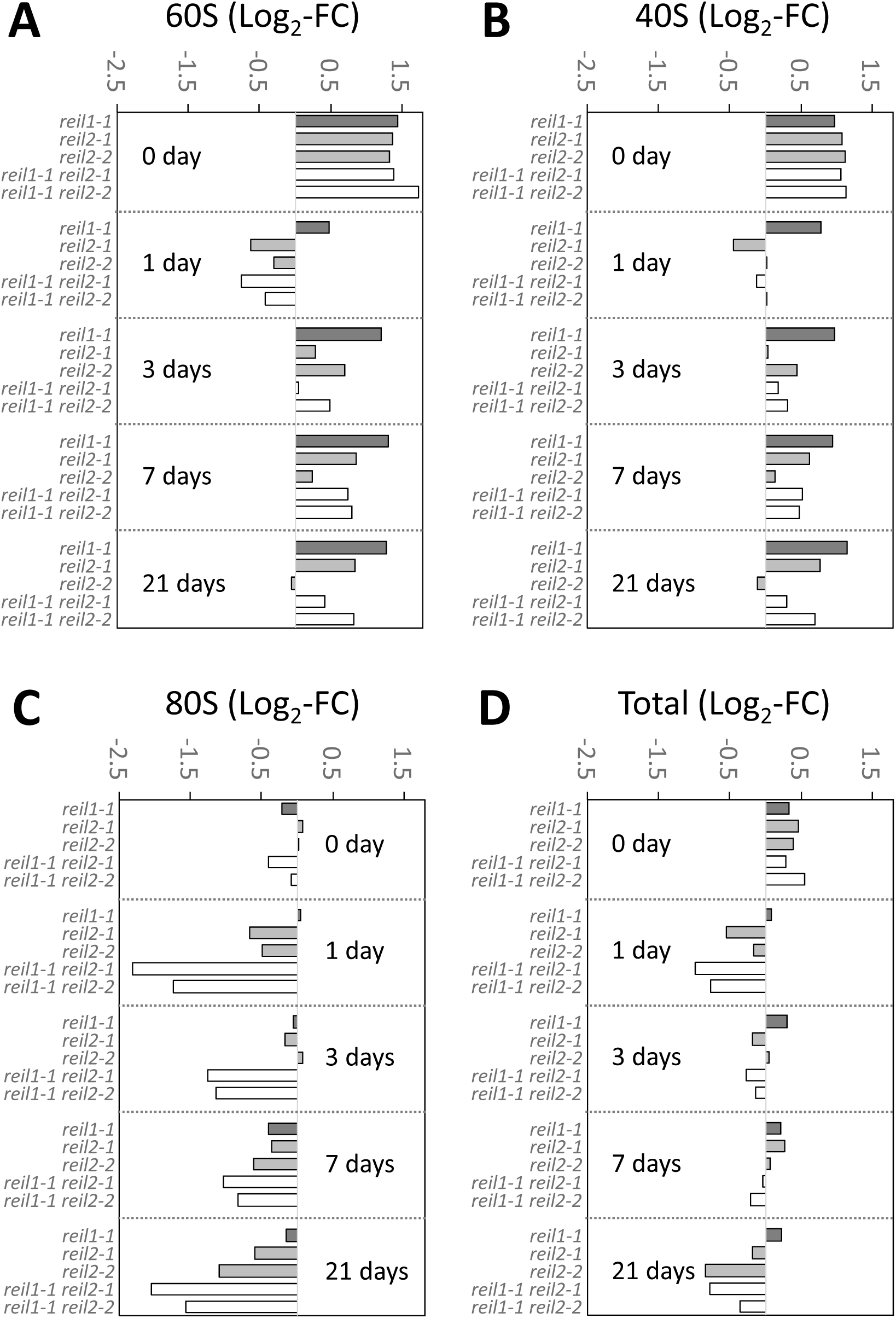
Relative quantification of abundances of ribosome complexes from non-acclimated (0 day) and 10°C cold acclimating *reil* mutants at 1, 3, 7, or 21 days after shift (cf. **Fig. 1**). Baseline corrected peak areas of ribosome complexes were integrated and log_2_-transformed ratios calculated relative to the respective wild type fractions at each time point, i.e. Log_2_-fold change (FC). The 60S LSU **(A)**, 40S SSU **(B)**, 80S monosomes **(C)** and the sum (“total”) of all ribosome complexes **(D)** were analyzed (dark grey*: reil1-1*, light grey: *reil2-1* and *reil2-2*, white: double mutants, *reil1-1 reil2-1* and *reil1-1 reil2-2*).

**Figure 3.**
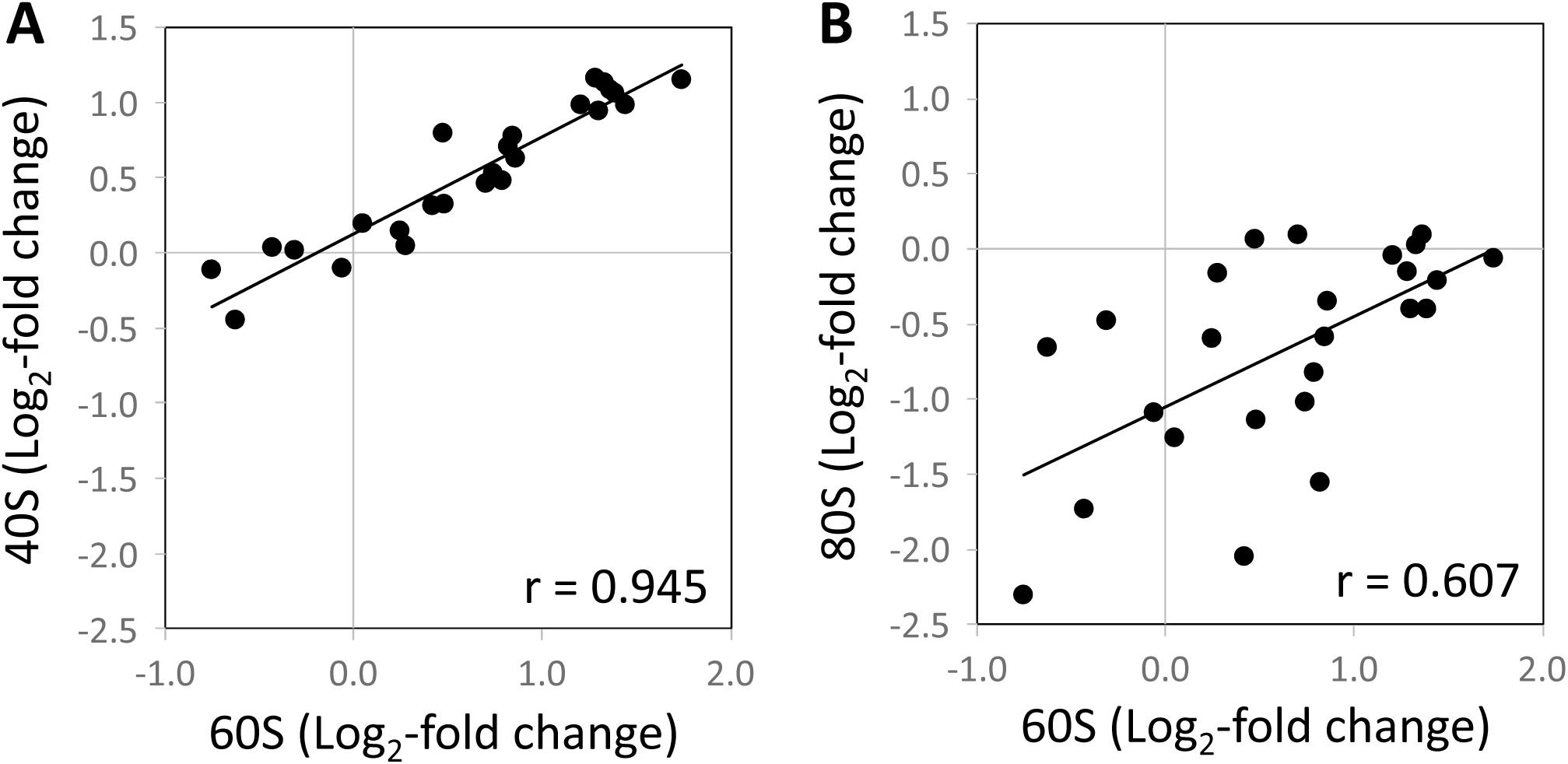
Correlation analyses of log_2_-transformed ratios of ribosome complex abundances comparing the cold induced changes of abundances relative to wild type of the 40S SSU **(A)** and of the 80S monosome fractions **(B)** to the changes of the 60S LSU fraction across all *reil* mutants (cf. **Fig. 2**). Inserts are the Pearson’s correlation coefficients (r) assuming a linear trend.

The double mutant effect on the 80S fraction was the largest and most consistent among the observed changes of cytosolic ribosome complexes. We exemplarily tested the 80S fractions of wild type and the *reil1-1 reil2-1* mutant for previously described heterogeneity (Martin and Hartwell, 1970; Zylber and Penman, 1970) using a sucrose sedimentation gradient tuned to separate translating ribosome fractions, specifically the 80S monosomes from the polysomes. Translating yeast or mammalian ribosomes are stable at high ionic strength. A sub-fraction of monosomes that are thought to be non-translating can be dissociated *in vitro* by elevating KCl levels during preparation (Martin and Hartwell, 1970; Zylber and Penman, 1970). Like in yeast and mammals, the 80S monosomes of plant roots partitioned into a KCl-sensitive and a stable fraction (**Fig. 4**). KCl concentrations that were elevated from regular 200 mM to 400 mM did not affect the root polysome fraction. In contrast, the 80S fraction that accumulated in Arabidopsis wild type roots at 21 days after cold shift was to a large extend KCl-sensitive as was indicated by associated decreases of 80S monosomes and increases of the 60S LSU and 40S SSU fractions (**Fig. 4A**). The 80S fraction of the *reil1-1 reil2-1* mutant contained only a minor amount of KCl-sensitive 80S monosomes (**Fig. 4B**). Like wild type, the polysome fraction of the *reil1-1 reil2-1* mutant was not KCl-sensitive. Polysome abundance was unchanged or slightly increased relative to wild type. We conclude that the failure of the double mutants to accumulate the 80S monosome fraction is likely a deficiency of accumulating the non-translating 80S sub-fraction. In summary, *reil* mutants appear to compensate deficiency of cytosolic ribosome biogenesis by over accumulation of free 60S LSU and 40S SSU and in the case of the double mutants by recruiting non-translating monosomes.

**Figure 4.**
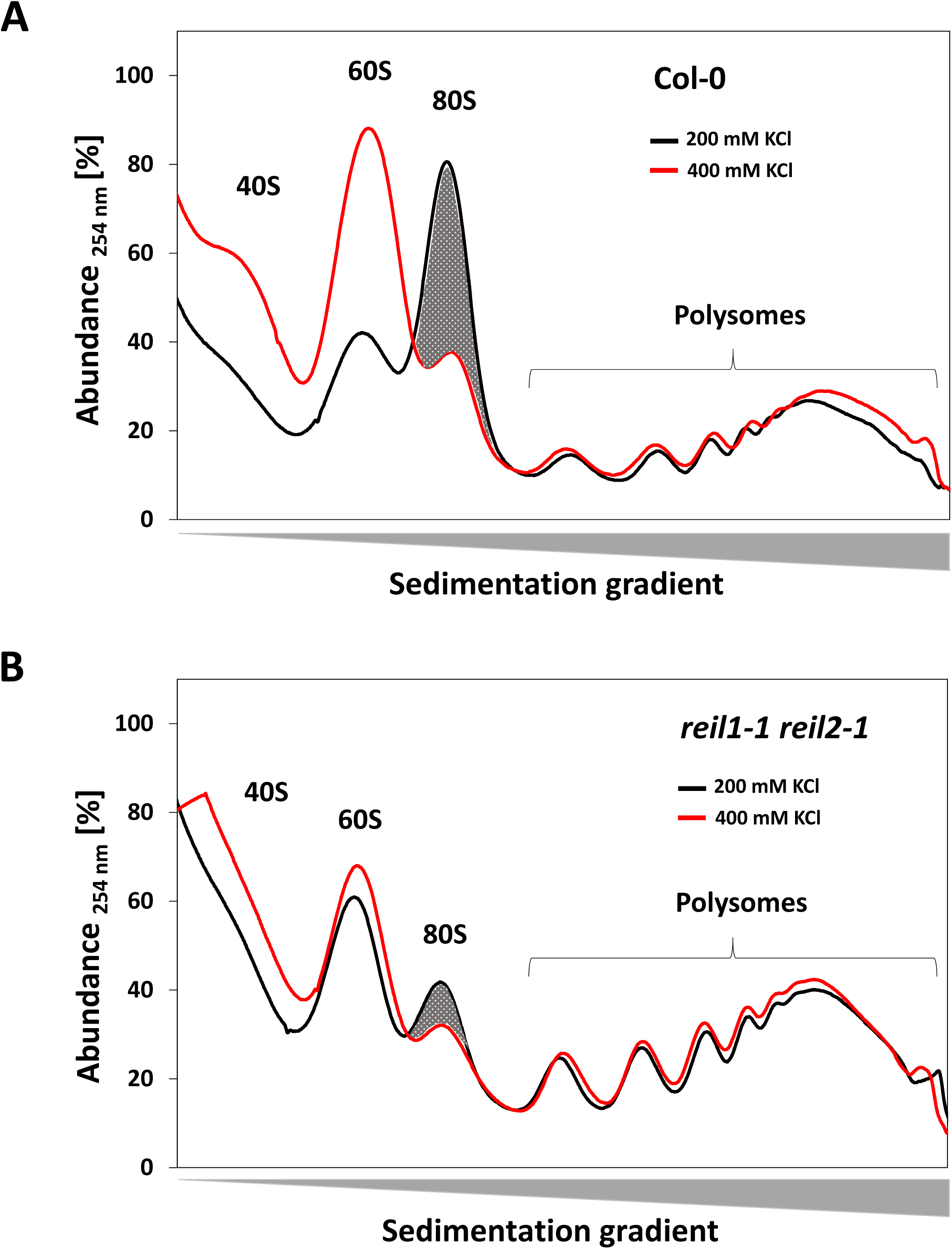
*In vitro* KCl-sensitivity test of monosomes and polysomes that were prepared from roots sampled at 21 days after start of 10°C cold-acclimation. Arabidopsis Col-0 wild type **(A)** was compared to the *reil1*-*1 reil2*-*1* double mutant **(B)**. Initial samples were homogenized and split into equal technical replicates. These replicates were extracted with PEB that either contained a normal KCl concentration (200 mM, black) or an elevated KCl concentration (400 mM, red). Note that the acclimated wild type accumulated a large fraction of KCl-sensitive monosomes, whereas KCl-sensitive monosomes were almost absent from cold acclimated *reil1*-*1 reil2*-*1* preparations.

### REIL deficiency deregulates transcriptional temperature acclimation responses in roots

In the following, we focused on the two double mutants, *reil1-1 reil2-1* and *reil1-1 reil2-2*, and compared non-acclimated (0 day) and the early cold-acclimating states at 1 day or at 7 days after cold-shift to Col-0 wild type. We chose the early cold-acclimating states according to our previous observations of a growth arrest after cold-shift that extended up to 7 day before Col-0 resumed growth in the cold (Beine-Golovchuk et al., 2018). Thereby, we aimed to avoid pleiotropic mutant effects that result from differential growth of the *reil* mutants at later time points. We aimed to reveal at transcriptional level, potential mechanisms that may regulate ribosome complex abundance or may take part in compensation responses to *reil* deficiency (**Supplemental Table S2**).

Col-0 and *reil* double mutant roots responded in our axenic, photomixotrophic, hydroponic growth system to 10°C cold shift by expected functional enrichment of transcript changes that relate to cold stress. Expression of genes belonging to the GOs, cold acclimation (GO:0009631) or cold response (GO:0009409), increased over time (**Fig. 5A**, **Supplemental Table S3**). Marker genes of cold acclimation and response, e.g. *CBF1*, *CBF2*, *CBF3*, *KIN1*, *KIN2*, *VIN3*, *COR15A* or *COR15B*, changed accordingly (**Supplemental Fig. S4**). Inversely to the cold response, expression of heat response genes (GO:0009408) significantly decreased (**Fig. 5A**). This observation differed compared to our previous experiments with soil cultivated rosette leaves (Fig. 8A of Beine-Golovchuk et al., 2018) and was accompanied by similar high light intensity (GO:0009644), hydrogen peroxide (GO:0042542) and reactive oxygen species (GO:0000302) responses. Together these observations indicated that the roots of Col-0 and those of the *reil* double mutants were stressed already at 20°C (**Fig. 5A**). This stress likely arose due to the artificial, light exposed hydroponic cultivation of roots in this study. Across the complete observation period, the *reil1-1 reil2-1* and *reil1-1 reil2-2* mutants shared constitutive reduction of expression of both *reil* genes in roots (**Supplemental Fig. S5**). The reduced gene expression in roots was similar to or in the case of residual *reil2* expression exceeded previous observations of *reil2* silencing in *reil1-1 reil2*-*1* leaves (Supplemental Table S2 of Beine-Golovchuk et al., 2018).

**Figure 5.**
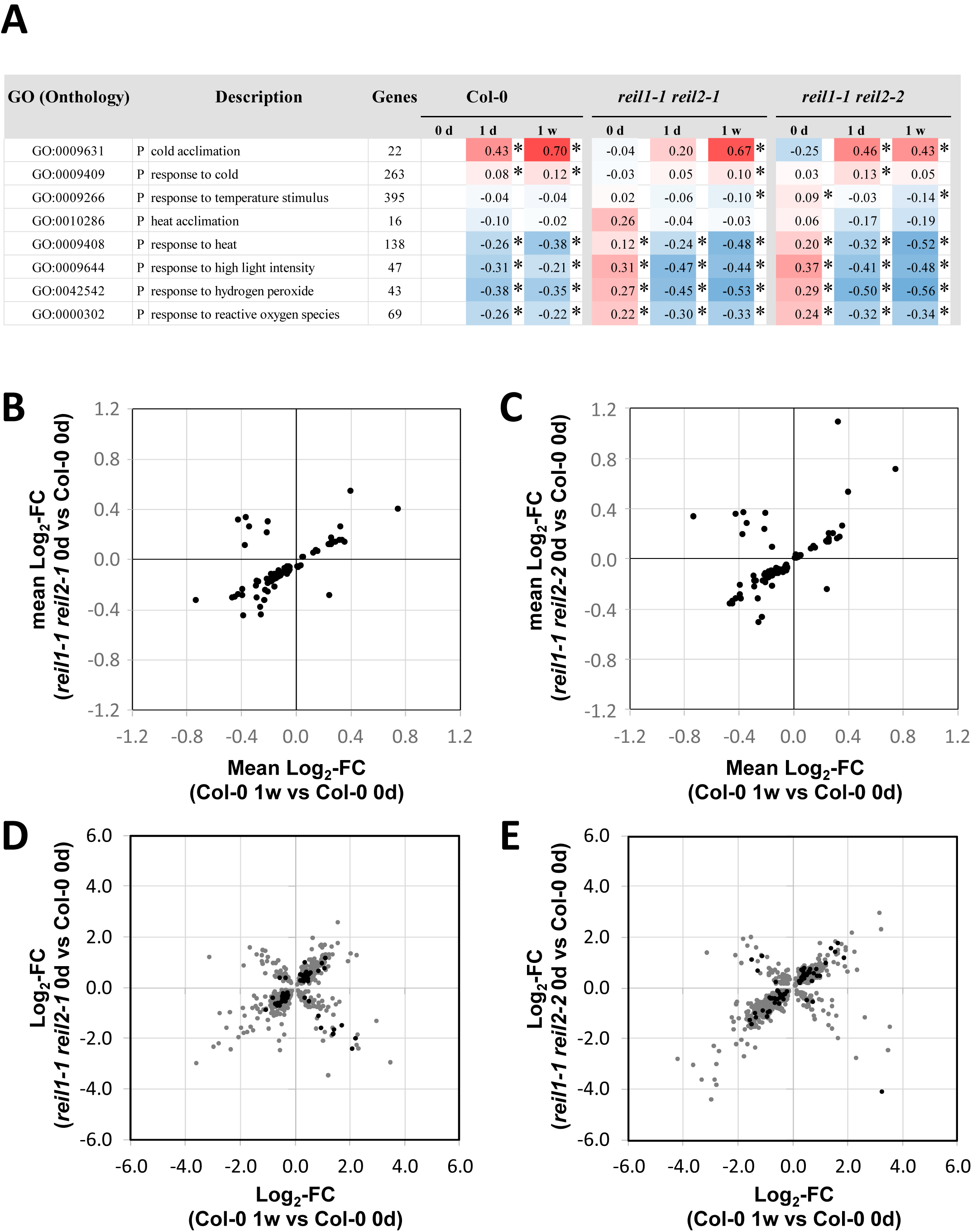
Functional enrichment and transcript correlation analyses of differential gene expression in the roots of Col-0, and the *reil1-1 reil2-1* and *reil1-1 reil2-2* double mutants at the non-acclimated state (0 day, 20°C) and shifted to 10°C cold for 1 day or 1 week. Differential gene expression was determined relative to non-acclimated Col-0 at optimized temperature 20°C. **(A)** Mean log_2_-fold changes (FC) of temperature related GO terms (P = biological process). Significant positive or negative functional enrichments, i.e. FDR-adjusted *P*-values < 0.05, are indicated by asterisks. The heat map color scale ranges from log_2_-FC +1.5 (red) to -1.5 (blue). Mean log_2_-FC, z-scores, and FDR-adjusted *P*-values of gene sets from 2145 GO terms were calculated by parametric analysis of gene set enrichment (PAGE; Tian et al., 2017; Supplemental Table S3). **(B)** Mean log_2_-FC of shared significantly enriched GO terms from the non-acclimated *reil1-1 reil2-1* mutant (0 d) compared to acclimated Col-0 at 1 week after shift. **(C)** Mean log_2_-FC of shared significantly enriched GO terms from the non-acclimated *reil1-1 reil2-2* mutant (0 d) compared to acclimated Col-0 at 1 week after shift. Note that both double mutants and cold acclimated Col-0 shared 100 GO terms of 2145 (FDR-adjusted *P* < 0.05). **(D)** Significant changed transcripts of non-acclimated *reil1-1 reil2-1* relative to non-acclimated Col-0 compared to significantly cold-responsive transcripts of Col-0 at 1 week after cold shift (P < 0.05, gray, P < 0.01 black). **(E)** Significant changed transcripts of non-acclimated *reil1-1 reil2-2* relative to non-acclimated Col-0 compared to significantly cold-responsive transcripts of Col-0 at 1 week after cold shift (P < 0.05, gray, P < 0.01 black). Note that correlation coefficients were not calculated because assumptions of linear correlation did not apply to the full set of observations.

Our previous study showed that mature leaves of *reil1-1 reil2-1* are prematurely cold acclimated at 20°C (Beine-Golovchuk et al., 2018). In our current study, we selected GOs that had significant enrichment of log_2_-FC in non-acclimated *reil* double mutant roots at 20°C compared to non-acclimated wild type at 20°C and of log_2_-FC in wild type that acclimated 7 days to 10°C compared to non-acclimated wild type at 20°C. We applied the threshold of FDR-adjusted *P* < 0.05 to each selection criterion (**Fig. 5 B-C**). The mean log_2_-FCs of genes from the selected GOs of the non-acclimated *reil* double mutants appeared to correlate to the mean log_2_-FCs of the cold acclimated wild type. However, part of the cold responsive GOs responded inversely (**Fig. 5 B-C**). The overlay of both, positively and negatively correlated gene expression in non-acclimated mutant roots compared to cold acclimated Col-0 wild type was even more obvious at single gene level (**Fig. 5 D-E**). In all of our four correlation analyses, namely the two double mutants at significant thresholds *P* < 0.05 and *P* < 0.01, respectively, the assumption of linear correlations did not apply. Therefore, correlation coefficients were not calculated. We concluded that *reil* double mutant roots deregulate transcriptional cold acclimation at 20°C in our growth system, with two subsets of genes, which either prematurely activate or prematurely deactivate. This response differs from premature cold acclimation as was indicated by transcription factors that control cold acclimation, e.g. *CBF1*, *CBF2* and *CBF3*. Transcripts of these factors did not consistently accumulate prematurely and expression of cold response marker genes, e.g. *KIN1*, *KIN2*, *VIN3*, *COR15A*, and *COR15B*, was reduced (**Supplemental Fig. S4**).

### REIL deficiency affects transcription of root morphogenesis genes and activates expression of cytosolic ribosome and translation related genes

Transcription of 34 GO terms had significant (FDR-adjusted *P* < 0.05) enrichment in both *reil* mutants at every time point of our study relative to non-acclimated Col-0 (**Supplemental Table S3**). These GO terms comprised the four previously mentioned stress stimuli (**Fig. 5A**) and two miscellaneous GOs, manganese ion binding, GO:0030145, and glycoside metabolic processes, GO:0016137 (**Supplemental Fig. S6**). A major mutant response pattern supported our observation of altered mutant root morphology (**Supplemental Fig. S3**). Twelve GO terms had constitutive reduced expression. These GOs were related to developmental processes, e.g. GO:0021700, root and root hair morphogenesis, e.g. GO:0010015 and GO:0010054, and cell wall organization and biogenesis, e.g. GO:0071554 (**Supplemental Fig. S6A**).

Eight of the remaining GO terms were related to translation or ribosomes, specifically to both subunits of cytosolic ribosomes (**Fig. 6A-B**). Col-0 transiently activated gene expression of cytosolic ribosomal proteins (RPs). *Reil1-1 reil2-1* and *reil1-1 reil2-2* consistently activated expression of cytosolic RP genes at 1day and 7 days after the cold shift. Expression of organelle RP genes did not significantly change in both mutants (**Fig. 6A**). The GO terms translation (GO:0006412) and the GO terms containing cytosolic RP genes (GO:0022625 - GO:0022627) responded similarly. Translation was specifically affected in translation initiation (GO:0003743) and eukaryotic translation initiation factor 3 (eIF3, GO:0005852) (**Fig. 6B**). The activation of cytosolic ribosome and translation related gene expression in mutant roots mirrored the effects in *reil1-1 reil2-1* leaves (Beine-Golovchuk et al., 2018). Kinetics of cold-induced gene expression appeared accelerated and reduced in amplitude in Col-0 roots compared to the rosette leaves analyzed in our previous study. In addition, expression of ribonucleoprotein (RNP) complexes (GO:0030529) and RNP biogenesis (GO:0022613) was activated in the cold. Expression of small nuclear RNP complexes (GO:0030532) that take part in splicing processes was constitutively activated in both mutants (**Fig. 6C**). Finally, five large biosynthesis and cytosol related GOs responded in a pattern that was similar to cytosolic ribosomes (**Supplemental Fig. S6B, Supplemental Table S3**).

**Figure 6.**
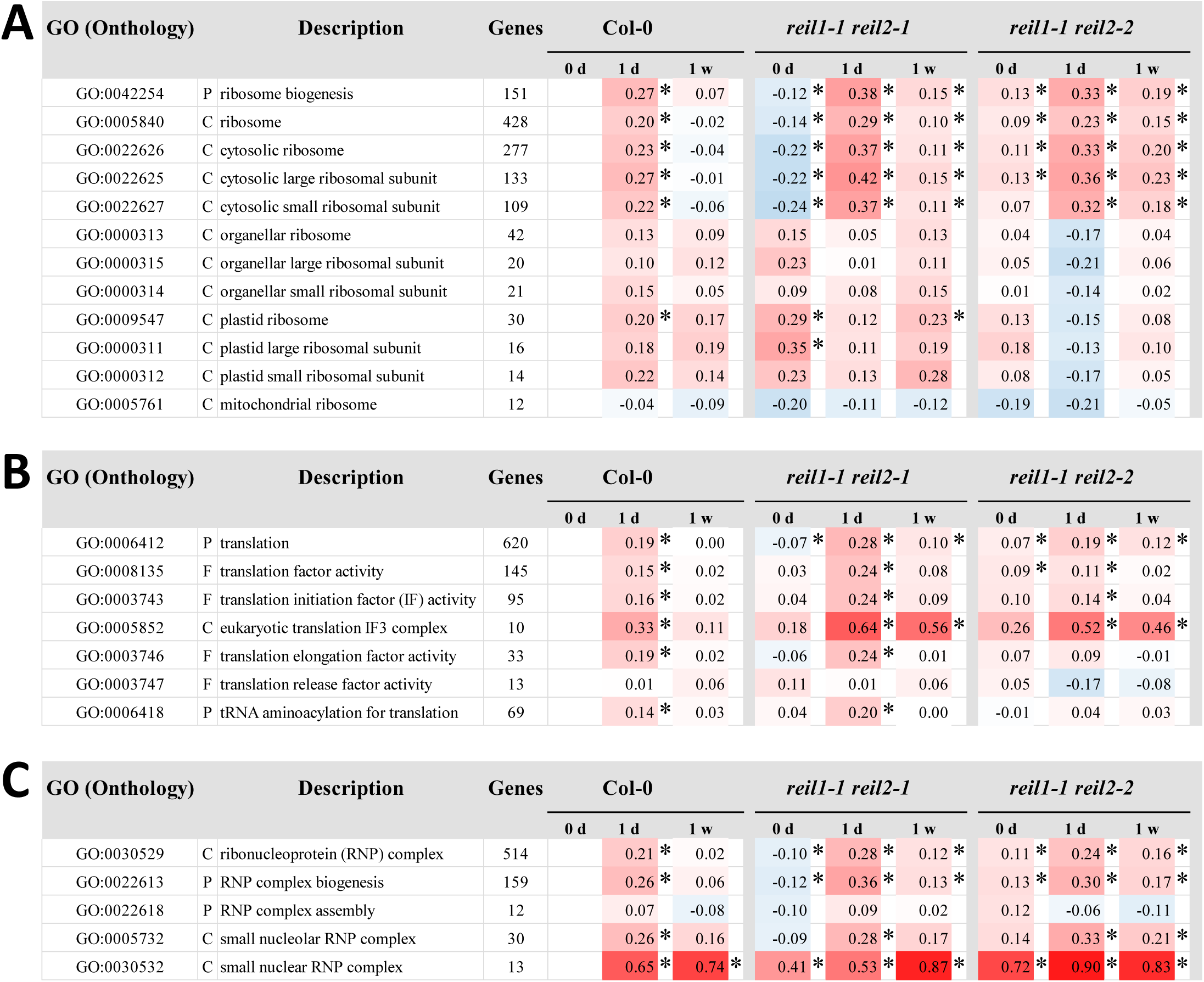
Functional enrichment analyses of differential gene expression in roots of Col-0, and the *reil1-1 reil2-1* and *reil1-1 reil2-2* double mutants at the non-acclimated state (0 day, 20°C) and shifted to 10°C cold for 1 day or 1 week. Differential gene expression was determined relative to non-acclimated Col-0 at optimized temperature 20°C. **(A)** Mean log_2_-fold changes (FC) of selected ribosome biogenesis and ribosome related GO terms. Note the prolonged and stronger activation of cytosolic ribosome related genes. **(B)** Mean log_2_-FCs of selected translation related GO terms. **(C)** Mean log_2_-FCs of selected ribonucleoprotein related GO terms. (C = cellular component, P = biological process, F = molecular function). Significant positive or negative functional enrichments, i.e. FDR-adjusted *P*-values < 0.05, are indicated by asterisks. The heat map color scale ranges from log_2_-FC +1.5 (red) to -1.5 (blue). Mean log_2_-FC, z-scores, and FDR-adjusted *P*-values of gene sets from 2145 GO terms were calculated by parametric analysis of gene set enrichment (PAGE; Tian et al., 2017). The full data set is listed in Supplemental Table S3.

A more detailed study of 40S SSU (104 transcripts covered by our current study) and 60S LSU (159 transcripts) RP gene expression compared to plastid 30S (30 transcripts) and 50S (41 transcripts) RPs revealed in part opposing effects of *reil* deficiency on structural cytosolic and plastid RPs (**Fig. 7**). At 10°C, expression of 40S and 60S RPs coordinately increased in both mutants relative to Col-0 at each time point. In the non-acclimated state, *reil1-1 reil2-1* roots reduced cytosolic RP expression whereas *reil1-1 reil2-2* roots had increased cytosolic RP transcripts (**Fig. 7A**). Expression of plastid RPs inversely increased in the non-acclimated state and decreased after shift to cold (**Fig. 7A**).

**Figure 7.**
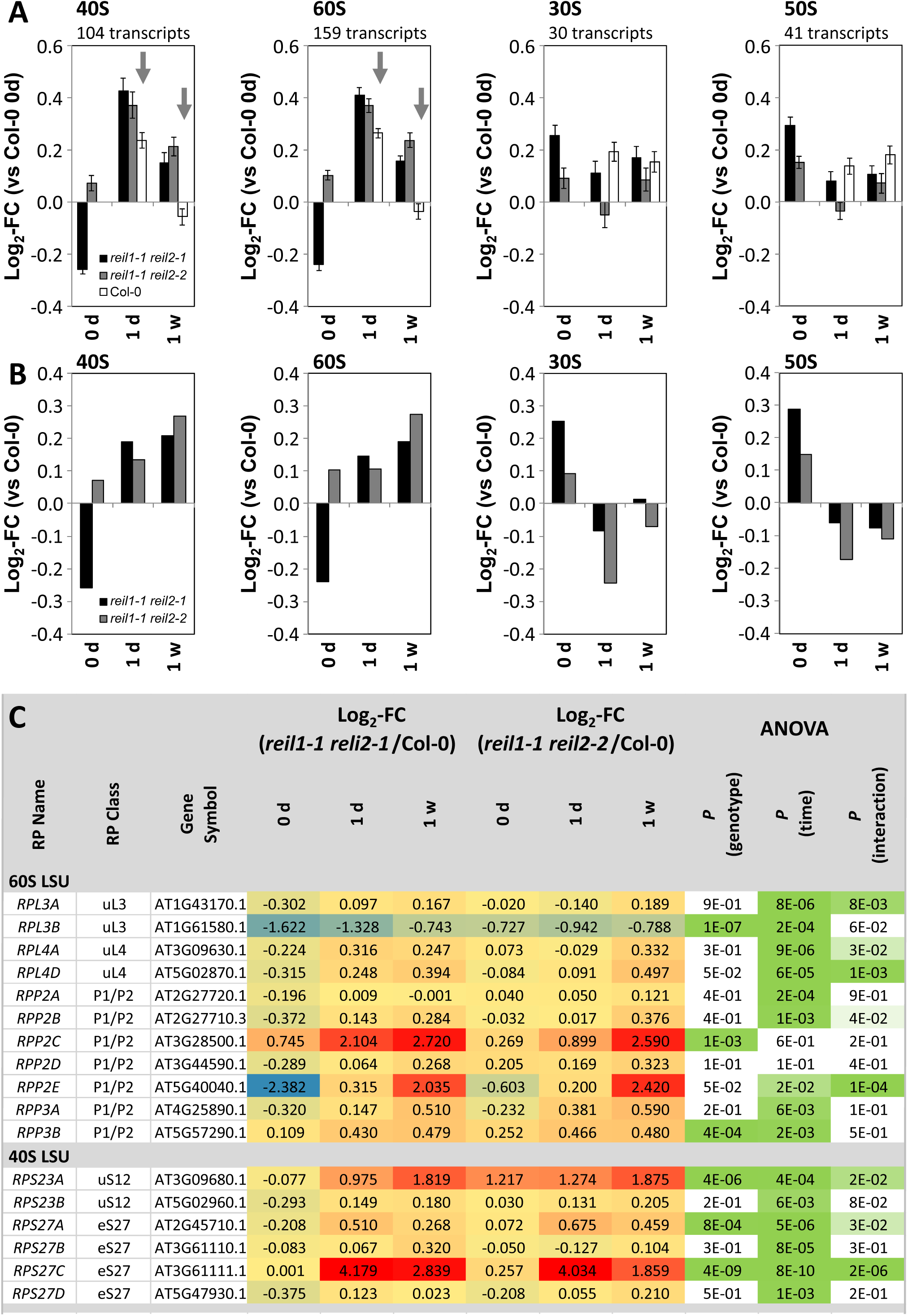
Differential expression of cytosolic and plastid ribosome genes in *reil1*-*1 reil2*-*1* and *reil1*-*1 reil2*-*2* double mutant roots at the non-acclimated state (0 day, 20°C) and shifted to 10°C cold for 1 day or 7 days. **(A)** Average log_2_-fold change (FC, means +/- standard error) relative to non-acclimated Col-0 (0 day) of transcripts coding for ribosome proteins (RPs) from the cytosolic 40S and 60S subunits and from the plastid 30S and 50S subunits. Grey arrows indicate transcript accumulation of the mutants in the cold. **(B)** Average log_2_-fold changes relative to Col-0 at each time point calculated form the means of the data from (A). **(C)** Log_2_-fold changes relative to Col-0 at each time point of selected cytosolic RP families. Two-factorial analysis of variance (ANOVA) indicates differential effects of the genotype, cold exposure (time) or the interaction of both on the expression of paralogous RPs from the 60S and the 40S subunits in the mutants. The three-color scale of the log_2_-FC heat map ranges from -3.0 (blue) to 0.0 (yellow) to ≥ +3.0 (red). The two-color significance scale ranges from *P* ≤ 1.0 x 10^-10^ (dark green) to *P* < 0.05 (light green), *P* ≥ 0.05 (white).

Analysis of cytosolic RP transcripts by pair wise comparisons at each time point and by two factorial analyses of variance (ANOVA) that tested for the influence of the factors, mutant genotype and time of cold exposure, indicated heterogeneity of RP gene expression at single gene levels (**Fig. 7B, Supplemental Table S2**). RP gene families comprised members with only a significant effect of either cold exposure or of genotype on their gene expression. We also observed RP paralogs with significant interactions between genotype and cold exposure or no significant change (**Fig. 7B**). In summary, *reil* deficiency globally activates cytosolic RP expression with inverse side effects in roots on plastid RP gene expression. *Reil* deficiency may affect gene expression of paralogs from one family synergistically, e.g. *RPL4A* and *RPL4D* (**Fig. 7B**). Alternately, the lack of REIL proteins can cause strong differential changes of gene expression of specific paralogs from 40S and 60S RP families, for example of the *RPL3*, *RPP2* or *RPP3*, and of the *RPS23*, and *RPS27* gene families (**Fig. 7B**). We conclude that differential regulation of RP gene transcription may contribute to the compensation of delayed cytosolic ribosome accumulation in the cold, but is likely not a compensation mechanism in the non-acclimated state.

### Constitutive transcriptional compensation responses to REIL deficiency indicate interaction of cytosolic ribosome maturation with nucleolar rRNA biogenesis and translation initiation

Several observations of GO enrichment among the differential transcript responses indicated constitutive transcriptional changes before and after cold acclimation, e.g. of the eIF3 complex (Fig. 6B). A detailed analysis of the ten members of GO:0005852 and the complete set of 13 subunits and in total 21 partly paralog genes constituting plant eIF3 (Burkes et al., 2001; Browning and Bailey-Serres, 2015) revealed a single gene, namely *eIF3C-2* (AT3G22860), that was highly and constitutively activated (**Fig. 8**). This activation was in contrast to its paralog, *eIF3C-1* (AT3G56150), and other eIF3 components, such as *eIF3G-2* (AT5G06000), that mostly mirrored cytosolic RP expression (**Fig. 8**). Applying the stringent criterion of significant constitutive changes relative to Col-0 in both mutants, at all non-acclimated and cold acclimating states, with *P* < 0.05 of each pairwise comparison (heteroscedastic T-tests), we discovered five additional constitutively accumulated transcripts. Next to two only marginally characterized F-box proteins (AT1G64540 and AT3G44120) and a MATH domain protein (AT3G29580) we found PISTILLATA (AT5G20240) and NUCLEOLIN 2 (NUC2, AT3G18610). PISTILLATA is a floral homeotic protein, and probable transcription factor involved in the control of petal and stamen development. PISTILLATA is in addition highly expressed in and near root quiescent centers (Lee et al., 2006; Brady et al. 2007). PISTILLATA appears to serve a yet elusive function in root development. Contrary to this enigmatic finding, function of NUC2 and of its paralog NUC1 provide a link to ribosome biogenesis (Durut et al., 2014). NUC1 and NUC2 nucleolins have antagonistic roles in rRNA gene expression. Both are required for plant growth (Sáez-Vásquez and Delseny, 2019). Similar to *eIF3C2* and *eIF3G2*, the *reil* double mutants constitutively activate *NUC2* expression whereas *NUC1* expression mirrored cytosolic RP gene expression. Contrary to NUC1, NUC2 maintains or induces a repressive state of rDNA chromatin (Durut et al., 2014).

In a meta-analysis, we compared constitutively changed root gene expression to non-acclimated and cold acclimating *reil1-1 reil2-1* rosette leaves at 1 day, 7 days, and 3 weeks after cold shift (Supplemental Table S2 of Beine-Golovchuk et al., 2018). Transcripts of *eIF3C2* (AT3G22860) and *NUC2* (AT3G18610) are constitutively and significantly increased in *reil1-1 reil2-1* mutant leaves with *P* < 0.05 (heteroscedastic T-tests) of each pairwise comparison to Col-0. Transcripts of the F-box proteins, AT1G64540 and AT3G44120, increase significantly using the same significance threshold in mutant leaves only at two or three of the compared time points, respectively. PISTILLATA and the MATH domain protein do not significantly change gene expression in *reil1-1 reil2-1* mutant leaves.

### Proteome Analysis of ribosome preparations from Arabidopsis root tissue

For comparative proteome analysis of ribosome complexes, we selected Col-0 wild type and the two double mutants, *reil1-1 reil2-1* and *reil1-1 reil2-2*. We performed two independent experiments each comparing pools of whole root systems from Col-0 wild type to pools of one of the two double mutants, DS1 (*reil1-1 reil2-2*) and DS2 (*reil1-1 reil2-1*). We analyzed the non-acclimated ribo-proteomes at day 0 to preparations obtained 7 days after shift to 10°C and sampled up to five non-translating ribosome complexes and the low-oligomer polysome fraction from sucrose density gradients. In this study, we avoided the period of reduced non-translating subunit abundance of the mutants at day 1 after cold shift and focused on the recovering ribo-proteome (**Fig. 1-2**).

We first analyzed the RP composition of each fraction to identify and align the ribosome fractions between experiments that may differ slightly in the position, but not the sequence of ribosome complexes with the sucrose gradient. For alignment of fractions, we used the sums of abundances of all detected 40S and 60S RPs, the plastid and mitochondrial 30S and 50S RPs, as well as the abundances of Arabidopsis EUKARYOTIC TRANSLATION INITIATION FACTOR 6, eIF6A (AT3G55620), and NONSENSE-MEDIATED mRNA DECAY 3 protein, NMD3 (AT2G03820) (**Supplemental Table S4**). eIF6A and NMD3 are homologs of the yeast cytosolic 60S maturation factors, TIF6 (Basu et al., 2001) and NMD3 (Ho et al., 2000). NMD3 and TIF6 are bound to translationally inactive pre-60S ribosome complexes (Lo et al., 2010; Greber et al., 2016) and in the case of TIF6 also indicate the position of NOP7-affinity purified 66S pre-ribosomes (Harnpicharnchai et al., 2001). Next to the polysome fraction, we obtained from both experiments a fraction enriched in 40S and organelle 30S RPs, designated 30S/40S fraction, a fraction of organelle 50S RPs (50S), a fraction that contained predominantly 60S RPs (60S) and a mixed fraction of 80S and 60S complexes (60S/80S) (**Supplemental Fig. S7**). In our current experiment, we did not reproducibly obtain proteome profiles of 80S fractions. For this reason, and because the 80S fraction is a difficult to interpret mixture of KCl-sensitive, non-translating and non KCl-sensitive, translating subpopulations (**Fig. 4**), we omitted 80S fractions from the current analyses. Both, the 60S and the 60S/80S fractions of experiments DS1 and DS2 contained eIF6A and NMD3 indicative of plant equivalents of 60S or 66S pre-ribosome complexes. In DS1, REIL proteins were detected only by few peptides and a sequence coverage of < 8.0 %. Therefore, we did not analyze the abundance of REIL proteins in DS1. In DS2, however, the 60S and 60S/80S fractions of the Col-0 wild type contained the REIL2 protein. Identification of REIL2 was supported by up to six peptides and 22.3 % sequence coverage.

The variation of cytosolic, plastid, and mitochondrial RP abundances among the four sample pools and resulting fractions of our proteomics experiments (**Supplemental Fig. S7**) prompted us to investigate total RP abundances added up across all ribosome fractions of each of the samples (**Supplemental Table S5**). The *reil1.1 reil2.1* sample of DS2 fully recovered after 7 days in the cold in terms of total 40S and 60S abundances. At the same time after cold shift, the *reil1.1 reil2.2* sample of DS1 still lacked in total 40S and 60S abundance. This observation was in agreement with our previous spectrophotometric analysis of the sedimentation profiles at seven days after cold shift (**Fig. 2 D**). The two mutant samples also differed in organelle RPs. *Reil1.1 reil2.1* roots decreased organelle RPs especially plastid and mitochondrial 50S RPs, both before and after cold shift. Inversely, *reil1.1 reil2.2* accumulated 50S RPs (**Supplemental Table S5**).

For our subsequent datamining, we focused on common responses of both mutants and searched for proteins that consistently accumulated or decreased relative to Col-0 wild type. We analyzed the non-translating fractions, namely the 60S and 60S/80S fractions in combination and separately the 30/40S fraction. Given the cold sensitivity of the *reil* mutants we looked for common changes of the two *reil* double mutants relative to Col-0 wild type in the cold and subsequently extracted the subset of common changes that were also present prior to the cold shift at 20°C. We decided to take into account the obvious variation 40S SSU and 60S LSU abundances among the ribosome fractions and among the four analyzed sample pools. To normalize the 60S and 60/80S fractions, we divided the abundance of the single observed proteins in each fraction to the abundance sum of all 60S RPs from the respective fraction. The 30S/40S fractions were normalized separately by the sum of all detected 40S RPs. The normalization enabled analyses of compositional changes of cytosolic RPs and co-purified non-ribosomal proteins in the non-translating SSU and LSU fractions. We analyzed increases and decreases in the mutants relative to the Col-0 wild type by log_2_-fold changes (log_2_-FC) (**Fig. 9-11**). The presence of a protein in a mutant sample and absence in corresponding Col-0 wild type rated as accumulation (+), absence in the mutant and presence in Col-0 as decrease (-).

**Figure 8.**
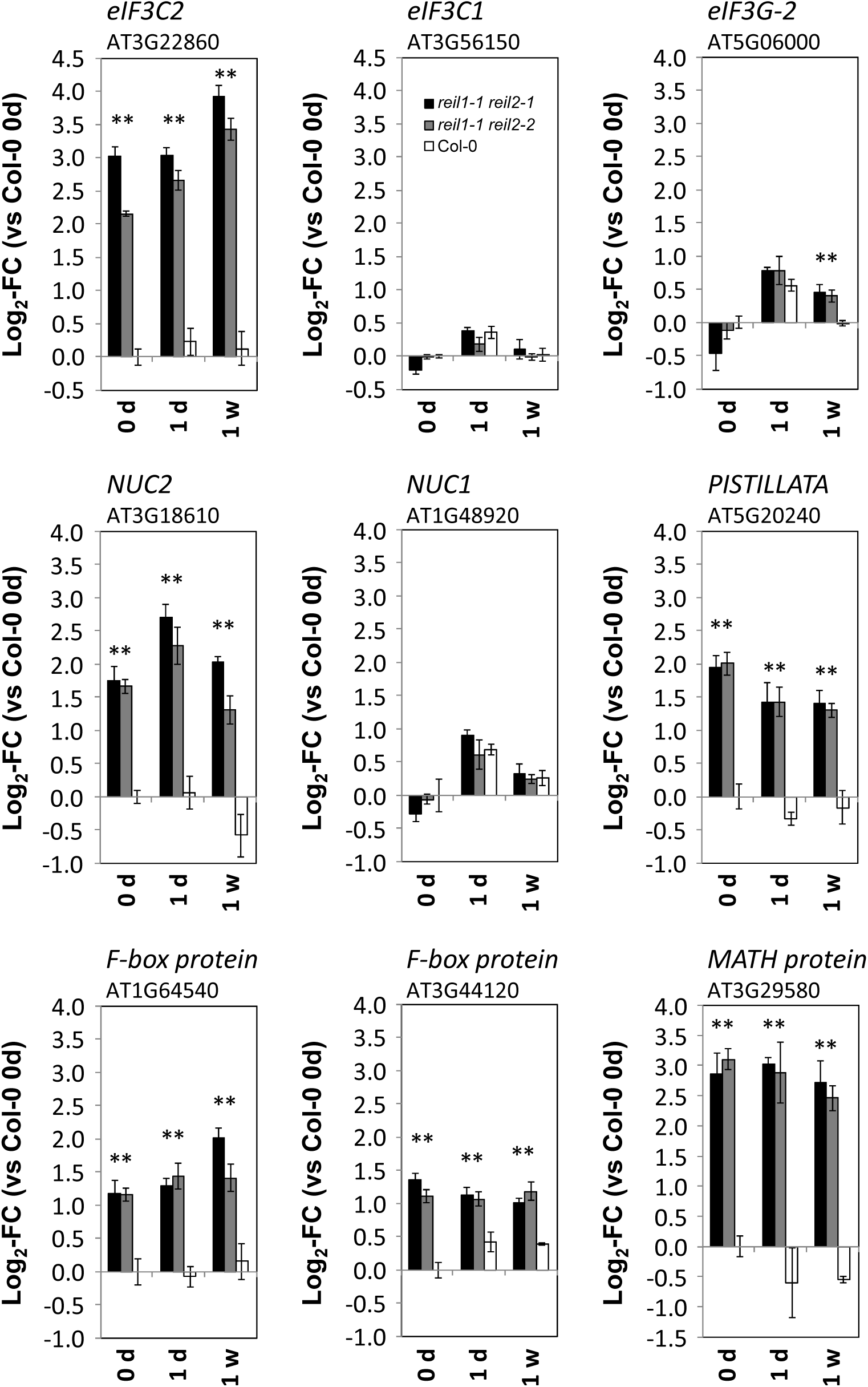
Differential expression of six genes that accumulate constitutively in *reil1-1 reil2-1* and *reil1-1 reil2-2* double mutant roots (means +/- standard error, n = 3). *eIF3C2*, *eIF3C1* and *eIF3G2* code for paralogs of the eukaryotic translation initiation factor 3 multi-protein complex. NUC1 and NUC2 are plant nucleolins. Asterisks indicate *P* < 0.05 (heteroscedastic Student’s t-test) of comparisons to Col-0 at the same time point. *NUC1*, *eIF3C1* and *eIF3G2* were added to this figure to demonstrate the specific mutant effect on paralog transcription.

**Figure 9.**
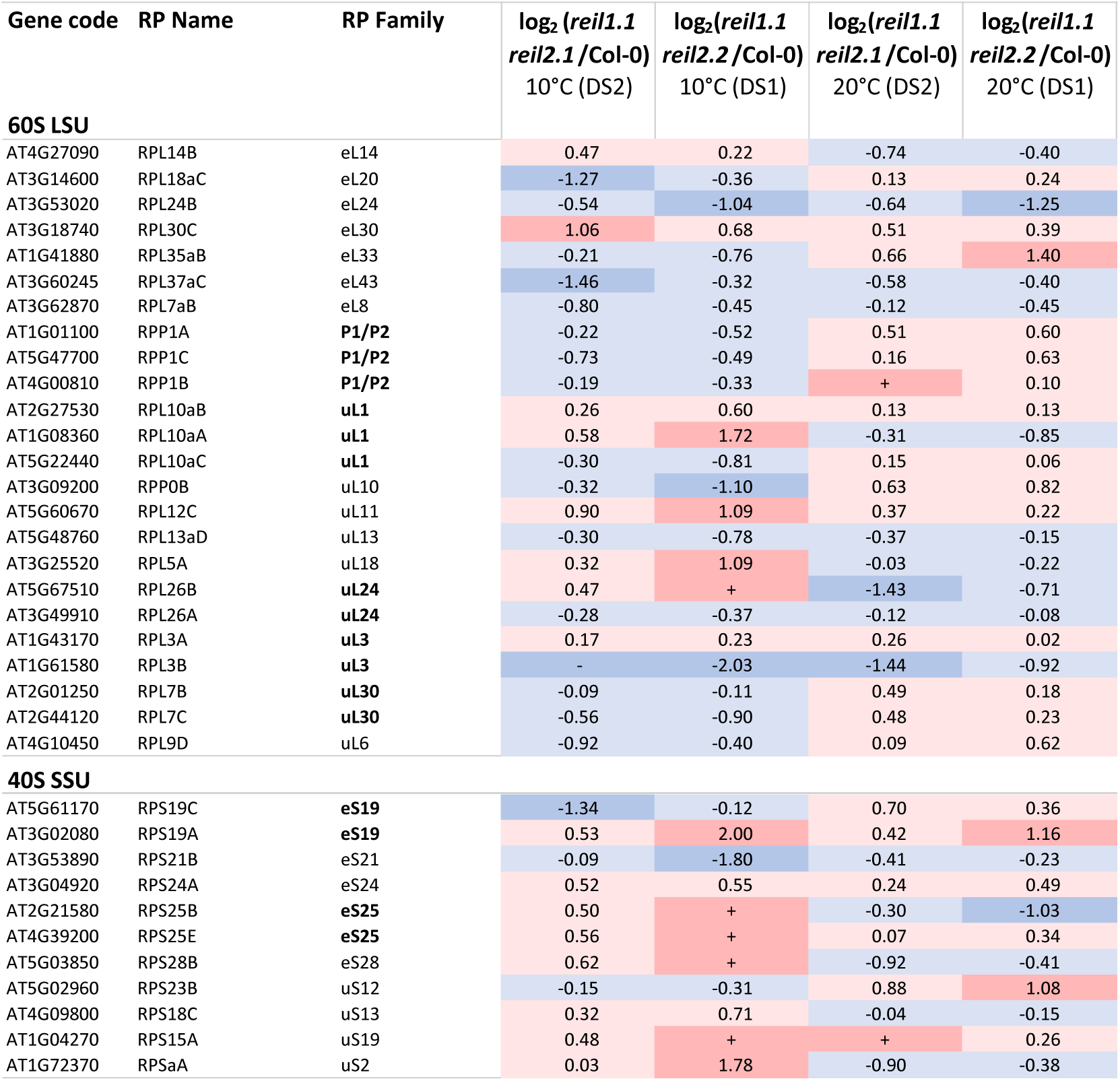
Changes of RP composition in the non-translating 60S LSU and 40S SSU fractions of the *reil1 reil2* double mutants 7 days after shift to 10°C and prior to cold shift at 20°C. This selection shows RPs with responses shared between *reil1-1 reil2-1* (DS2) and *reil1-1 reil2-2* (DS1) at 10°C and 20°. Log_2_-fold changes between mutants and Col-0 wild type were calculated after normalization of RP LFQ-abundances by the abundance sums of all detected 60S RPs in the respective fraction and combined across the 60S and 60S/80S fractions (top) or by the abundance sums of all detected 40S RPs in the 30S/40S fractions (bottom). Presence relative to absence in Col-0 and log_2_-fold increases > 1 are color-coded red, increases < 1 are coded light-red. Absence relative to presence in Col-0 and log_2_-fold decreases < -1 are color-coded blue, decreases > -1 are coded light-blue.

As to general observations, DS1 and DS2 differed in complexity. DS1 yielded a more complex data set with 2820 compared to 1675 detected proteins in DS2 (**Supplemental Table S5**). The number of detected cytosolic RPs and RP paralogs was, however, almost equal, 175 in DS1 and 174 in DS2 (**Supplemental Table S6**). The experiments shared 162 cytosolic RPs and RP paralogs of in total 1485 common proteins. The detection of cytosolic RPs was mostly paralog specific or represented RP families with identical amino acid sequences. Twenty-six (DS1) and 21 (DS2) cytosolic RPs lacked paralog specificity (**Supplemental Table S6**). Our analysis did not differentiate splice variants. In agreement with the cold-sensitivity of the mutants, we found more shared changes of normalized protein abundances in the cold than prior to cold shift. Most of the observed changes were larger in magnitude in DS1 (*reil1.1 reil2.2*) than in DS2 (*reil1.1 reil2.1*). The current experiments, however, do not allow attributing these observations to differences between the two mutants or to a potential slight difference between their respective recovery kinetics after cold shift (**Fig. 2**).

### 60S RPs and RP paralogs differentially accumulate in non-translating 60S fractions of *reil* double mutants

We selected 61 proteins with shared increases (36) or decreases (25) at 10°C in the non-translating 60S fractions of *reil* double mutants (**Supplemental Table S7A**). Co-purified proteins with an abundance maximum in the 50S fraction were removed from our selection of 60S compositional changes. Most observed changes were below 2-fold. Eight proteins increased consistently at 10°C and 20°C, seven decreased. The remaining 46 proteins changed in part inversely at 20°C in both *reil* double mutants (15), were not detectable at optimized temperature or differed at optimized temperature between mutants (**Supplemental Table S7A**).

The majority of the selected proteins were structural cytosolic RPs (45) that either increased (20) or decreased (25) relative to the sum of abundances of all detected 60S RPs. RPL3A (AT1G43170), RPL10aB (AT2G27530), RPL12C (AT5G60670), and RPL30C (AT3G18740) increased consistently in the *reil* double mutants, RPL3B (AT1G61580), RPL7aB (AT3G62870), RPL13aD (AT5G48760), RPL24B (AT3G53020), RPL26A (AT3G49910), RPL37aC (AT3G60245) decreased (**Fig. 9**). The 61 changes comprised cytosolic RPs that are assembled in the nucleus and RPs that are added or exchanged within the cytosol, such as, already mentioned RPL24B (AT3G53020) of the eL24 family and RPL10A (AT1G14320) and RPL10C (AT3G11250) of the uL16 family, RPP0B (AT3G09200) and RPP0C (AT3G11250) of the uL10 P-stalk family or RPP1A (AT1G01100), RPP1B (AT4G00810), RPP1C (AT5G47700), RPP2D (AT3G44590) of the P1/P2 P-stalk protein family (**Fig. 9, Supplemental Figure S8, Supplemental Table S7A**). RPP1A, RPP1B, RPP1C and RPP0B decreased at 10°C and increased at 20°C relative to wild type (**Fig. 9, Supplemental Figure S8**).

### Homologs of yeast 66S pre-ribosome constituents and 60S biogenesis factors accumulate in non-translating 60S fractions of *reil* double mutants

The sixteen non-structural cytosolic RPs among the set of 61 differentially accumulating proteins were with two exceptions, namely a root tip expressed LEA protein (AT5G60530) and a R3H domain protein (AT1G03250), homologs of yeast 60S biogenesis factors and 66S pre-ribosome constituents (**Fig. 10**). We found the homologs of yeast TIF6, NMD3, RLP24, NOP7, NOG1, NOG2 (alias NUG2), RPF2, NSA2, MAK16, RSA4, CIC1 (alias NSA3), EBP2, HAS1 and YTM1. All accumulated in the non-translating 60S fractions at 10°C. The TIF6-, NMD3- and RLP24-homologs and the R3H domain protein accumulated in addition at 20°C. Next to yeast NOP7, the biogenesis factors, yeast TIF6, NOG1, MAK16, HAS1, EBP2, NSA2, CIC1, YTM1 and RLP24 are non-ribosomal proteins that are associated with 66S pre-ribosomes that can be NOP7 affinity-purified (Harnpicharnchai et al., 2001; Kater et al., 2017). The remaining NMD3, NOG2, RPF2, RSA4 are part of other pre-60S ribosome complexes. In detail and again assuming basic homology of plant and yeast 60S biogenesis, NMD3 is a nuclear export adaptor of pre-60S subunits that is part of a pre-60S ribosome carrying TIF6, LSG1 and the yeast REI-homolog, REH1 (Ma et al., 2017). Like TIF6, RLP24 and NOG1, NMD3 is released from pre-60S ribosomes in the final cytosolic maturation steps and recycled (Ho et al., 2000, Greber et al., 2016). NOG2 (alias NUG2) precedes NMD3 binding to pre-60S ribosomes at overlapping binding sites prior to nuclear export (Matsuo et al., 2014). RSA4 is present on NOG2 associated particles and co-substrate of the REA1 remodeling factor that is required for NOG2 release from pre-60S ribosomes (Matsuo et al., 2014). Finally, NOG2 (Gamalinda et al., 2013) associates with pre-60S ribosomes already in the nucleolus and together with NSA2 is thought to be part of a 66SB pre-ribosome and required for 27SB pre-rRNA processing into 25S rRNA (Gamalinda et al., 2013). Similarly, RFP2 (Morita et al., 2002) predominantly localizes to the nucleolus, associates with 66S pre-ribosomes by an alternative recruiting pathway and is also required for 27SB pre-rRNA processing (Gamalinda et al., 2013). HAS1 is RNA helicase that takes part in both small and large ribosomal subunit biogenesis (Gnanasundram et al., 2019). HAS1 is present in yeast 90S pre-ribosomes and has multiple binding sites in the 18S rRNA and the 5.8S and 25S rRNAs and remains associated with pre-40S and pre-60S complexes (Gnanasundram et al., 2019). We, however, detected the plant homolog of HAS1 only in non-translating 60S fractions of the cold-exposed *reil* double mutants (**Supplemental Table S7A**).

**Figure 10.**
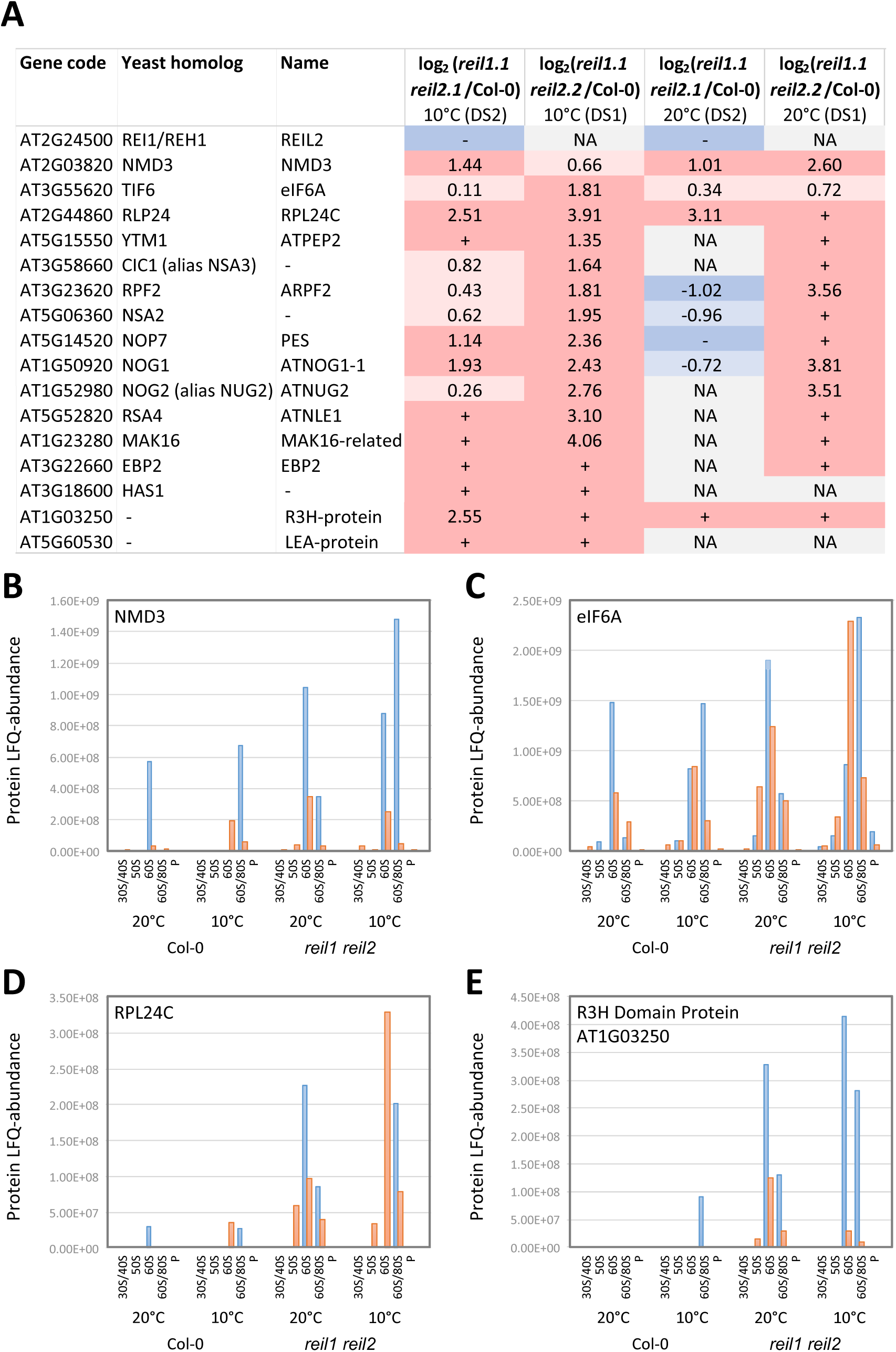
Changes of the composition of ribosome associated proteins in the non-translating 60S fractions of the *reil1 reil2* double mutants 7 days after shift to 10°C and prior to cold shift at 20°C. **(A)** REIL2 and the shared changes at 10°C between *reil1-1 reil2-1* (DS2) and *reil1-1 reil2-2* (DS1). Log_2_-fold changes between mutants and Col-0 wild type were calculated after normalization of protein LFQ-abundances by abundance sums of all detected 60S RPs and combined across the 60S and 60S/80S fractions. Presence in mutants relative to absence in Col-0 and log_2_-fold increases > 1 are color-coded red, increases < 1 are light red. Absence relative to presence in Col-0 and log_2_-fold decreases < -1 are color-coded blue, decreases > -1 are light-blue. Absence in both mutant and Col-0 is color-coded grey and indicated by NA (not available). **(B, C, D, E)** LFQ-abundance distributions of NMD3, eIF6A, RPL24C, and R3H domain Protein AT1G03250 across the sampled ribosome fractions and analyzed conditions, DS1 orange, DS2 blue. Note the abundance maxima of the proteins in the non-translating 60S and 60/80S fractions.

**Figure 11.**
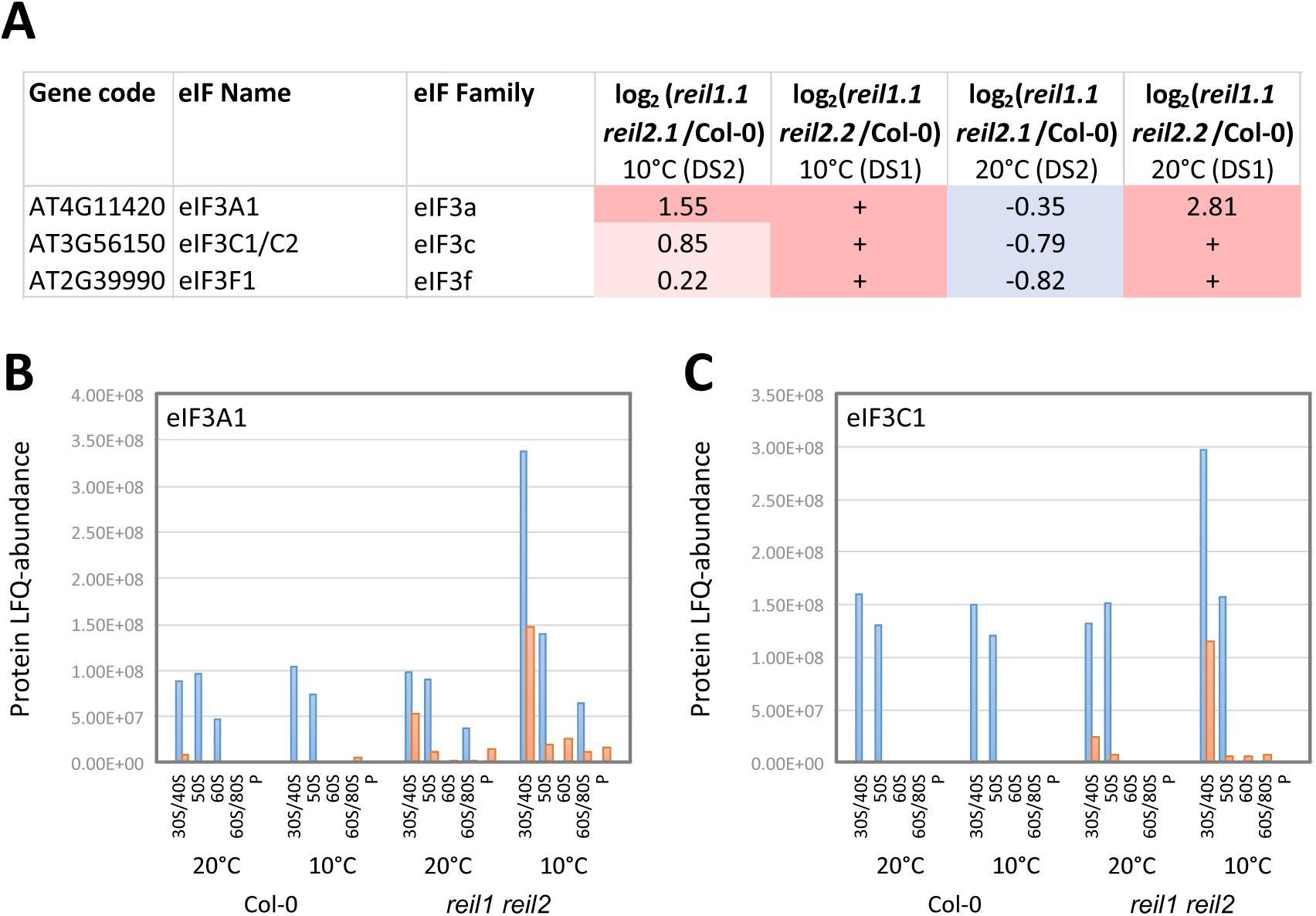
Changes of the composition of translation initiation factor 3 proteins in the non-translating 40S fraction of the *reil1 reil2* double mutants 7 days after shift to 10°C and prior to cold shift at 20°C. **(A)** Shared changes at 10°C between *reil1-1 reil2-1* (DS2) and *reil1-1 reil2-2* (DS1). Log_2_-fold changes between mutants and Col-0 wild type in the 30S/40S fractions were calculated after normalization of protein LFQ-abundances by abundance sums of all detected 40S RPs. Presence in mutants relative to absence in Col-0 and log_2_-fold increases > 1 are color-coded red, increases < 1 are light red. Absence relative to presence in Col-0 and log_2_-fold decreases < -1 are color-coded blue, decreases > -1 are light-blue. **(B)** Distribution of eIF3A1 LFQ-abundances across the sampled ribosome fractions and analyzed conditions, DS1 orange, DS2 blue. **(C)** Distribution of eIF3C1 LFQ-abundances across the sampled ribosome fractions and analyzed conditions, DS1 orange, DS2 blue. Note the abundance maxima in the 30/40S fractions that may contain 40S subunits and 43S pre-initiation complexes.

### 40S RPs and eukaryotic translation initiation factors differentially accumulate in non-translating 40S fractions of *reil* double mutants

In comparison to the moderate number of highly specific ribosome and ribosome associated proteins (RAPs) that differentially accumulated in non-translating 60S fractions, we detected 146 proteins with shared increases (135) or decreases (11) at 10°C in the non-translating 40S fractions of the *reil* double mutants (**Supplemental Table S7B**). Only 21 of the selected proteins were structural cytosolic RPs that either increased (12) or decreased (9) relative to the sum of abundances of detected 40S RPs. The majority of selected proteins were co-purified proteins, e.g. 2 organelle 30S RPs, 3 nitrilases, 4 heat shock proteins, 6 components of the proteasome, 10 components of ATP synthases and a multitude of enzymes. We concluded that these observations may be relevant, but are likely not linked to the non-translating 40S fraction. RPS24A (AT3G04920), RPS19A (AT3G02080), RPS25E (AT4G39200), and RPS15A (AT1G04270) increased consistently in the *reil* double mutants, RPS21B (AT3G53890) decreased (**Fig. 9**). Six additional 40S RPs changed inversely. Besides changes of structural 40S RPs, we found, at 10°C, accumulation of eIF3A1 (AT4G11420), eIF3C1 (AT3G56150), and eIF3F1 (AT2G39990) in the non-translating 40S fraction of both double mutants (**Fig. 11**). Detection of eIF3C1 (AT3G56150) included minority contribution by two of 16 (DS1) or one of 22 peptides (DS2) of eIF3C2 (At3g22860) (**Supplemental Table S6**). These selected translation initiation factors are a subset of the ten detected components of the 13-subunit plant eukaryotic translation initiation factor 3 (eIF3) (Browning et al., 2001). eIF3 is part of the 43S and 48S pre-initiation complexes and exits upon joining of the 60S subunit (Browning and Bailey-Serres, 2015).

### Association of differential accumulation of RPs and RAPs in non-translating 40S and 60S fractions of *reil* double mutants with differential gene expression

With few exceptions, the differential accumulation of RPs in non-translating 40S and 60S fractions and the changes of ribosome biogenesis factors or eIF3 components in these fractions did not correlate with respective changes in transcript levels. We attempted to correlate differential accumulation of RPs in non-translating 40S and 60S fractions of *reil* double mutants from experiments DS1 and DS2 with differential accumulation of their transcripts at 20°C prior to cold shift and at 7 days after shift to 10°C (**Supplemental Figure S9 A-B, E-F**). In addition, we tested for time-shifted correlation of differential protein accumulation at 7 days after cold shift with preceding differential gene expression at 1 day after the shift (**Supplemental Figure S9 C-D**). With few exceptions, differential protein accumulation was not generally associated with changes of gene expression. However, the consistent decrease of *RPL3B* (AT1G61580) gene expression (**Fig. 7C**) before and after cold shift, average log_2_-FC = -1.03, *P* = 1.15 10^-7^ (genotype effect of 2-way ANOVA), matched to consistent reduction of RPL3B protein in the non-translating 60S fraction of the two *reil* double mutants, experiments DS1 and DS2 (**Fig. 9**). In addition, consistent slight increase of *RPL24C* (AT2G44860) gene expression, average log_2_-FC = 0.24, *P* = 1.43 10^-2^ (genotype effect of 2-way ANOVA), matched to consistent increase of RPL24C protein in the non-translating 60S fraction of the two *reil* double mutants.

## Discussion

### The mixotrophic *in vitro* cultivation system: requirements and biases

We validated through this study the function of Arabidopsis REIL proteins as ribosome biogenesis factors of eukaryotic plant ribosomes. Our previous studies of REIL function in rosette leaves (Beine-Golovchuk et al., 2018) indicated that analyses of eukaryote ribosome complexes from leaf material were in part masked by almost equally abundant chloroplast ribosomes. Using root material, the organelle interference of UV-absorbance traces from density gradient separations of plant ribosome complexes was negligible (**Fig. 1**). Plastid and mitochondrial ribosomes co-purified from root material, but the prokaryote-type ribosome complexes were according to our proteome analyses on average more than 100-fold less abundant than eukaryote ribosome complexes (**Supplemental Figure S7**, **Supplemental Table S5**). The choice of root material thus enabled relative and correlative quantification of eukaryote ribosome complexes (**Fig. 2-3**). These analyses were not possible previously using leaf material (Beine-Golovchuk et al., 2018). Analyses of ribosomes complexes required amounts of ∼100 mg (FW) tissue (Beine-Golovchuk et al., 2018). Our axenic *in vitro* cultivation system (**Supplemental Figure S1**) delivered required amounts of root tissue from plants that had developmental stage, ∼1.10 (Boyes et al., 2001). Our current study confirmed *reil* gene expression in Col-0 wild type roots under our mixotrophic hydroponic cultivation conditions and strong reduction of expression of both genes in *reil* double mutants (**Supplemental Figure S5**). Recent promoter GUS studies confirmed constitutive *reil2* expression throughout Arabidopsis seedlings under similar *in vitro* conditions and revealed relevance of REIL2 in the root organ by demonstrating high expression in the root tip (Yu et al., 2020).

The previously reported delay of *reil* mutant development and growth in the cold attenuated using our cultivation system (**Supplemental Figure S2-S3**). We did not adjust temperature settings to achieve previous phenotypic strength. Instead, we used the phenotype-attenuating settings to minimize pleiotropic effects that may arise due to differential development of mutants and wild type (**Supplemental Figure S2**). We maintained comparability with our previous analyses of soil grown Arabidopsis rosettes (Schmidt et al., 2013; Beine-Golovchuk et al., 2018) by selecting stage ∼1.10 plants to perform temperature shift experiments.

We did not succeed, however, in avoiding the bias of mixotrophic cultivation. Mutant plants stayed dwarfed and flowered early even at optimized temperature without addition of sucrose to the liquid cultivation medium. As consequence, we cultivated in the presence of sucrose, but need to consider that the sucrose supply will mask or distort endogenous carbohydrate dependent signals. In addition, the illumination of the root system and the high humidity of hydroponic cultivation may affect transcriptional responses, for example, the observations linked to high light and reactive oxygen species that we report (**Fig. 5A**).

We discovered a further potential developmental bias of our study. *Reil* mutations affected root architecture (**Supplemental Fig. S3**). The branching pattern of hydroponic root systems in *reil* single and *reil1 reil2* double mutants appeared changed. The functional link of REIL proteins to root development or root system architecture (Jung and McCouch, 2013) was supported further by functional enrichment analyses of differential gene expression. The *reil1 reil2* double mutants had altered transcript levels of genes belonging to developmental growth ontologies, specifically root morphogenesis, epidermis and trichoblast differentiation (**Supplemental Figure S6**). These transcript changes between mutants and wild type became apparent already at optimized cultivation conditions and preceded cold induced responses. Because our current cultivation system was not suited for detailed root morphology or root patterning studies, we did not pursue this aspect further and considered this effect negligible for our current study.

We previously reported premature transcriptional cold acclimation responses at optimized temperature of *reil1.1 reil2.1* rosette leaves from soil-cultivated plants (Beine-Golovchuk et al., 2018). An initial global differential gene expression analysis of shared significantly enriched GOs between the non-acclimated *reil1 reil2* mutants and 7-days cold acclimated Col-0 appeared to confirm the triggering of premature transcriptional cold responses (**Fig. 5B-C**). The same comparison at single gene level, however, revealed an overlay of positively and negatively correlated transcriptional responses (**Fig. 5D-E**). Globally, premature responses to heat among the 138 genes of this ontology appeared to dominate over the cold responses of 263 genes, respectively, but neither cold acclimation nor heat acclimation responses were significantly enriched in the *reil1 reil2* double mutants prior to cold shift (**Fig. 5A**). Selected marker genes of cold acclimation and cold response were either not prematurely changed, e.g. *C-REPEAT/DRE BINDING FACTOR1/ DEHYDRATION-RESPONSIVE ELEMENT BINDING PROTEIN1B* (*CBF1/DREB1B*), *CBF2/DREB1C*, *CBF3/DREB1A*, and *VERNALIZATION INSENSITIVE 3* (*VIN3*), or had reduced expression prior to cold shift, e.g. *COLD-INDUCED1* (*KIN1*) and *KIN2* or *COLD-REGULATED15A* (*COR15A*) and *COR15BA* (**Supplemental Figure S4**). Cold acclimation of Arabidopsis has a well characterized carbohydrate component, where acclimating plants accumulate, e.g. (Kaplan et al., 2004; Kaplan et al., 2007) and deacclimating plants rapidly reduce endogenous sugar levels (Pagter et al., 2017). Adaptations in photosynthetic carbon metabolism provide signals that are integral part of Arabidopsis cold acclimation, including post transcriptional and transcriptional changes to sucrose metabolism (Stitt and Hurry, 2002). In addition, photosynthesis-derived glucose is a regulator of TARGET OF RAPAMYCIN (TOR) signaling that is a component of Arabidopsis cold acclimation, e.g. (Xiong et al., 2013; Dong et al., 2019). Considering the biases of our root cultivation system and especially the masking of carbohydrate signals, we cautiously conclude that the *reil1 reil2* double mutants deregulate temperature responses in mixotrophic roots at optimized temperature.

### REIL deficiency reveals requirement for *de novo* synthesis of eukaryote ribosomes after cold shift

Our current study confirmed and extended our previous observation that *reil* mutants delay accumulation of non-translating 60S LSUs after cold shift (Beine-Golovchuk et al., 2018). We demonstrated that the delay affects both, the non-translating 60S LSUs and 40S SSUs (**Fig. 1-2**). Shortage of non-translating free ribosome subunits was clearly transient and the balance of the abundances of non-translating large and small eukaryotic ribosome subunits was under tight control in plants (**Fig. 3**). We currently do not have evidence that control of this balance is dependent on REIL function in Arabidopsis. Our observation of concomitant decrease of 40S SSUs in *reil* double mutants was unexpected, as the 40S SSU fraction increases in yeast Δ*rei1* mutants, e.g. (Lebreton et al., 2006; Greber et al., 2016). Arabidopsis apparently has different control mechanisms to balance non-translating 40S and 60S subunits. However, our finding agrees with yeast data that show concomitant decreases of 40S SSUs and 60S LSUs in mutants that are defective in 60S structural RPs, whereas mutants defective in 40S structural RPs may accumulate high levels of 60S LSUs (Cheng et al. 2019).

We observed compensation responses of REIL deficiency that act both, before cold shift and after extended periods of cold (**Fig. 2-4**). All *reil* mutants over-accumulated baseline pools of non-translating 60S LSUs and 40S SSUs, when not immediately responding to cold shift. Mutants containing a defective *reil2* gene depleted in addition the 80S fraction in the cold. In Arabidopsis, the 80S fraction, similar to yeast (Martin and Hartwell, 1970; Zylber and Penman, 1970), appears to be composed of translating and non-translating 80S ribosome monomers. Our KCl-sensitivity tests of the Arabidopsis 80S fraction and the poor correlation between 80S and non-translating 60S fractions indicated the heterogeneous nature of the 80S fraction (**Fig. 3-4**). Mainly for this reason, we did not analyze the ribo-proteome of the 80S fraction in this study.

The function of the two REIL paralogs was clearly not fully redundant. REIL1 deficiency can be compensated successfully under both optimized and low temperature conditions, but a compensation by over-accumulation of free subunits and slight deficiency after cold shift was observable (**Fig. 2A-D**). In contrast, REIL2 deficiency was compensated at optimized temperature, but compensation apparently failed in the cold. This was indicated by persistent depletion of the 80S fraction both in the *reil2* single mutants and in the *reil1 reil2* double mutants (**Fig. 2C**).

The requirement of *de novo* synthesis in the cold is evident by slow recovery of ribosome complex abundances in the *reil* mutants. The time scale of 3-7 days after the shift is in agreement with previous reports of ribosome protein turnover in Arabidopsis (Li et al. 2017; Salih et al., 2019), where half-life of the ribosome population was approximately 3-4 days. Only RPP0D and the RACK1B and 1C paralogs were shorter lived with half-lives between 0.5 and 1.5 days (Salih et al., 2019). Induction of 40S and 60S RP transcripts after cold shift in the Col-0 wild type and compensatory enhanced and prolonged expression of these genes in *reil1 reil2* double mutants indicate that Arabidopsis roots (**Fig. 6, 7A, 7B**) are prepared by transcript availability for ribosome *de novo* synthesis after cold shift. Rosette leaves provided similar observations, however, at larger amplitude (Beine-Golovchuk et al., 2018).

Compensational depletion of non-translating ribosome complexes in the *reil* mutants cannot be explained alone by 60S subunit shortage and deficient *de novo* ribosome biosynthesis. We propose that Arabidopsis maintains and controls pools of non-translating free 60S LSUs and 40S SSUs, likely under a tight and common control (**Fig. 3**) and in addition a pool of assembled, but non-translating 80S ribosomes. These translationally inactive pools of ribosome complexes recycle ribosome complexes and may serve as buffers that respond to changing translation demands (Uesono and Toh-e, 2002; Krokowski et al., 2011; van den Elzen et al., 2014). Such pools may save energy by harboring surplus extant ribosomes instead of initiating wasteful premature degradation. Such pools may fill, when translation demands are low and prepare for rapid responses to environmental cues without immediate need of comparatively slow *de novo* ribosome biosynthesis and assembly. Fluctuating temperature cues, such as investigated in this study, obviously alter translation demands.

### Effects of REIL deficiency on translation initiation

In yeast cells, Rei1 deficiency, similar to other mutations that limit the amount of 60S LSU subunits, is associated to the hallmark observation of so-called half-mer polysomes, e.g. (Parnell and Bass, 2009; Greber et al., 2016). Half-mers are polysome complexes with a stalled 40S preinitiation complex and one or more fully assembled translating 80S ribosomes bound to a single mRNA. Stalling of the preinitiation complex finds an obvious explanation through the lack of translationally competent 60S LSUs for translating 80S monomer assembly. Half-mer polysomes of yeast have been detected by sucrose density gradient centrifugation (Parnell and Bass, 2009; Greber et al., 2016). In our sucrose gradient analyses, we found indications of accumulating half-mer polysomes in the *reil1 reil2* double mutants at 1-3 days after cold shift (**Fig. 1**), but this phenomenon did not persist after prolonged cold exposure (**Fig. 1, 4**). Proteomic analysis demonstrated the presence of components of the eIF3 multi-protein complex (Burks et al., 2001) in the non-translating 40S fraction and in a subsequent fraction enriched for organelle 50S RPs. These observations likely indicate the presence of co-purified plant 43S and 48S preinitiation complexes, respectively (**Fig. 11B, C**). Accumulation of eIF3 components in both cold acclimating *reil1 reil2* double mutants indicated accumulation mostly of stalled 43S preinitiation complexes after cold shift (**Fig. 11A-C**). The existence of half-mer polysome complexes in cold exposed *reil* double mutants was supported by the presence of eIF3A1 in the mutant low-oligomer polysome fraction in one of our analyses (**Fig. 11B**). We conclude that REIL protein deficiency can cause accumulation of Arabidopsis 43S preinitiation complexes and that half-mer polysomes can accumulate. These observations prove REIL function as a ribosome biogenesis factor by indicating lack of translationally competent 60S subunits in the cold.

At transcriptome level, we found evidence that *reil* deficiency feeds back constitutively onto gene expression of the *eIF3C2* paralog. REIL deficiency specifically released the suppression *eIF3C2* expression (**Fig. 8**) and largely did not affect expression of other constituents of the eIF3 complex. In our current study, the eIF3C2 protein was not selectively detectable with one of 22 or two of 16 detected peptides shared between the eIF3C1 and eIF3C2 paralogs (**Supplemental Table S6**). The 805 amino acid eIF3C2 protein has an internally shortened N-terminal domain and lacks a large part of the C-terminus compared to the 900 amino acid eIF3C1 paralog (Burks et al., 2001). It may be allowed to speculate that the eIF3C2 paralog competes with eIF3C1 in plant eIF3 complexes and/ or may have an inhibitory regulatory role that adjusts eIF3 abundance to availability of translationally competent 60S LSUs. Such a function may cause a sub-fraction of eIF3 not to bind to ribosomes, but a mechanistic explanation must currently remain elusive.

### Effects of REIL deficiency on ribosome biogenesis

The *reil1 reil2* mutants accumulate Arabidopsis homologs of yeast 60S and 66S ribosome biogenesis factors in the non-translating 60S fraction. Homologs of cytosolic biogenesis factors, NMD3, TIF6, RLP24 that are assembled within the nucleus and released in the cytosol before the 60S LSU becomes translationally competent accumulate before and after temperature shift (**Fig. 10A-E**). Nuclear ribosome biogenesis factors associate predominantly with the 66S pre-ribosome. These factors include proteins that block positions in the 66S complex which are later occupied by the cytosolic biogenesis factors in the 60S pre-ribosome. Nuclear ribosome biogenesis factors accumulated in the cold in both *reil* double mutants and only in one double mutant, *reil1.1 reil2.2* before cold shift (**Fig. 10A**). These observations indicate a heterogeneity between the *reil2* alleles that we currently have available or a non-controlled variation in our cultivation system at optimized temperature that currently eludes us.

We therefore conclude that REIL deficiency inhibits the release of Arabidopsis NMD3-, TIF6-, and RLP24-homologs and thereby slows down biogenesis of translationally competent 60S LSUs. We interpret accumulation of nuclear ribosome biogenesis factors as pile-up of precursor complexes caused by blocked cytosolic 60S maturation. We observe that the block of 60S maturation intensifies at low temperature (**Fig. 10**). Taken together the observations prove conserved REIL function as a cytosolic 60S biogenesis factors. In addition, we imply previously non-described plant proteins, namely a R3H-domain protein and a LEA-like protein that do not have apparent homology in yeast or humans beyond the conserved R3H-domain, to contribute to plant specific aspects of 60S biogenesis.

At transcriptome level, we find *NUC2*, but not *NUC1* gene expression constitutively activated. *NUC2* codes for one of the two antagonistic plant nucleolins and thereby provide a functional link to eukaryote ribosome biogenesis. Both, *NUC2* and *NUC1* expression is required for plant growth. NUC2 and NUC1 act as histone chaperons early in eukaryotic ribosome biogenesis. The nucleolins control rDNA variant expression by modulation of 45S rRNA transcription and/ or processing (Durut et al., 2014; Sáez-Vásquez and Delseny, 2019). *NUC2* expression is high in the root apical meristem, similar to our still enigmatic finding of *reil* dependent PISTILLATA deregulation in roots. Both nucleolins are expressed in root and leaf tissues, but *NUC2* transcripts accumulate more in root than in leaf and shoot tissue (Durut et al., 2014). Durut and co-authors (2014) conclude that NUC2 might bind nucleosomes to induce and/or maintain a repressive rDNA chromatin state. Building on the hypothesis that *NUC2* may act as a repressor (Sáez-Vásquez and Delseny, 2019), we may naively interpret our observation of deregulated *NUC2* expression as indication of feedback repression of the primary step of rRNA transcription in response to blocked 60S maturation. However, studies on *reil2* single paralog mutants suggest that the control of rRNA transcription may be negligible compared to feed back control of rRNA processing exerted downstream of the 35S rRNA precursor (Yu et al., 2020). Obviously, this control affected both of the two parallel terminal plant pre-rRNA processing pathways (Weis et al., 2015). Accumulation of pre-60S maturation factors confirmed stalling of nuclear 66S maturation complexes that contain partially processed rRNA.

### REIL deficiency affects the RP paralog composition of non-translating 40S SSU and 60S LSU fractions

We discovered that REIL deficiency affects the paralog composition of non-translating ribosome subunits. Most of our observations are less than 2-fold changes (**Fig. 9**). Considering the composite nature of the 40S SSU and 60S LSU fractions that may contain both, recycled ribosome populations that were synthesized prior to cold shift and ribosome populations that are *de novo* synthesized in the cold, we were surprised by these observations that were consistent between the two *reil* double mutants.

One of the more obvious observations was the consistent decrease of RPL24B of the eL24 RP family in the non-translating 60S fractions of the *reil1 reil2* double mutants (**Fig. 9**). This phenomenon can be linked directly to the concomitant accumulation of the RPL24C (**Fig. 10**), the Arabidopsis homolog of the yeast RLP24 maturation factor that is a placeholder during pre-60S complex assembly and replaced in the cytosol to form translationally competent 60S LSUs (Greber et al., 2016). Competent 60S LSUs of yeast contain either yeast RPL24A or RPL24B. Similar to yeast, Arabidopsis REIL proteins appear to be required to facilitate or accelerate this exchange.

Two other 60S RP families are assembled onto the 60S subunit in the cytosol, uL10 and uL16 (Greber et al., 2016). Of these, we find RPP0B, uL10 family, to be consistently decreased at 10°C and inversely increased at 20°C in both *reil* double mutants (**Fig. 9**). RPP0B is one of the Arabidopsis paralogs that code for the P-stalk protein P0 that anchors the P-stalk to the ribosome (Krokowski et al., 2005; Krokowski et al., 2006; Mitroshin et al., 2016; Liljas and Sanyal, 2018). P0 interacts as a scaffold with other P-stalk proteins of the P1/P2 family, here represented by consistent abundance changes of Arabidopsis RPP1A, RPP1B and RPP1C (**Fig. 9**) and likely overall reduced abundance of RPP2D in the *reil* double mutants (**Supplemental Figure S8A**). The fully assembled and active P-stalk substructure interacts with translation elongation factors and is required for translation.

Besides these changes of ribosome subunit composition that may be explained by lack of direct functions of the REIL biogenesis factors in the cytosol, we found temperature dependent and independent effects on the composition of structural ribosome proteins within the non-translating 40S SSU and 60S LSU (**Fig. 9**). Such composition changes must involve either an indirect REIL function that affects *de novo* subunit assembly in the nucleus or may involve RP paralog exchange in assembled ribosome that pre-exist in the cytosol. Such a mechanism was previously proposed for *in situ* ribosome repair (Holt and Schuman, 2013). Among the many observations of compositional changes, we would like to highlight the effect on the uL3 family that is part of 60S LSUs (Kim et al., 1990). REIL proteins apparently are required to accumulate RPL3B paralogs in non-translating 60S fractions (**Fig. 9**). The slight accumulation of abundant RPL3A, an essential gene for Arabidopsis (Meinke, 2019), appears to compensate reduction of RPL3B in our study. In addition, RPL3B decrease in the absence of REIL proteins has a transcriptional component, as the reduction of *RPL3B* transcript is one of the rare observations of otherwise absent associations between transcriptome and proteome changes of non-translating fractions (**Supplemental Figure S9**).

## Conclusion

Our study proves that Arabidopsis REIL proteins function as cytosolic ribosome biogenesis factors and supports their role of accelerating ribosome *de novo* synthesis in the cold. Arabidopsis compensates for the lack of REIL function at optimized temperature, but cannot fully compensate in the cold. The compensation mechanisms accumulate non-translating pools of ribosome complexes and appear to activate pre-existing pools upon cold-shift. We hypothesize, that Arabidopsis controls non-translating ribosome complexes to buffer fluctuating demands on translation. Thereby Arabidopsis may limit the demand for *de novo* ribosome synthesis under rapidly changing environmental conditions. REIL function is mostly post-transcriptional, however, with a few obvious transcriptional components. REIL function feeds forward onto formation of 43S initiation complexes likely in a passive mode by modulating 60S LSU availability or more actively via *eIF3C2*, *NUC2* or RP gene expression and feedback onto nuclear pre-60S subunit assembly. Through disturbance of *de novo* ribosome biosynthesis and translation initiation, *reil* mutants allow the study of these processes and their control in plants.

Unexpectedly, REIL function in Arabidopsis appears to control the paralog composition of non-translating 40S and 60S subunits. Considering paralog composition as one mechanism that may create heterogeneous functional ribosome populations (Xue and Barna, 2012; Simsek et al., 2017; Shi et al., 2017; Genuth and Barna, 2018), we propose that *reil* mutants and temperature acclimation cues may provide feasible and highly relevant experimental systems to study ribosome heterogeneity in plants.

## Supplementary data

**Supplemental Figure S1.** Hydroponic growth system of *Arabidopsis thaliana* Col-0 wild type and mutants. The system was feasible for temperature shift experiments in the vegetative phase and for rapid sampling of pooled material of whole root or shoot systems.

**(A)** Assembled growth system with stainless steel mesh adjusted level with the surface of ∼ 250 mL liquid Murashige and Skoog-medium with 2% sucrose. One of the four agar blocks for seed germination is indicated (red arrow, view from the side).

**(B)** Assembled growth system with stainless steel mesh adjusted level with the surface of the liquid. The four agar blocks for seed germination are indicated (red arrows, view from the top). Each block contains a single seed.

**(C)** Assembled growth system with *Arabidopsis thaliana* Col-0 wild type plants at approximate ten-leaf stage after 4 weeks in 16 h/ 8 h long day conditions at 20°C during the day and 18°C during the night (view from the side).

**(D)** Assembled growth system with *Arabidopsis thaliana* Col-0 wild type plants at approximately ten-leaf stage (view from the top).

The clamped-on glass lids were opened and sterile conditions compromised for the photographs.

**Supplemental Figure S2.** Exemplary documentation of shoot systems of hydroponically cultivated *Arabidopsis thaliana* wild type (Col-0), of the single mutants, *reil1-1, reil2-1, reil2-2,* and of the double mutants, *reil1-1 reil2-1 (DKO1)* and *reil1-1 reil2-2 (DKO2),* before (0 day) and 7 or 21 days after shift from 20°C (day)/ 18°C (night) to 10°C (day) and 8°C (night).

Note that at 21 days after cold shift shoots and inflorescences of *reil2-1, reil2-2,* and the double mutants, *reil1-1 reil2-1 (DKO1)* and *reil1-1 reil2-2 (DKO2),* are smaller than wild type and the *reil1-1* mutant. Growth of the double mutants did not arrest under the conditions of this hydroponic system. Cultivation was in liquid MS media with 2% sucrose (w/v) adjusted to pH 5.7 (Murashige and Skoog, 1962). All photographs were taken separately as indicated by vertical white bars. In parts, black background was added to the single sections of the graph for a regular and centered display of the shoot systems. All bars are 1 cm and indicate the slightly varying scales of single photographs.

**Supplemental Figure S3.** Exemplary documentation of hydroponic root systems of *Arabidopsis thaliana* wild type (Col-0), of the single mutants, *reil1-1*, *reil2-1*, *reil2-2*, and of the double mutants, *reil1-1 reil2-1* (*DKO1*) and *reil1-1 reil2-2* (*DKO2*), before (0 day) and 7 or 21 days after shift from 20°C (day)/ 18°C (night) to 10°C (day) and 8°C (night).

Roots were cut at the hypocotyl to root transition. Root systems of single plants were carefully prepared with minimal wounding from joined cultivations of four plants in single containers. Preparation of complete root systems from single plants was not possible. Photographs may therefore show in parts incomplete root systems. Note the shortened primary root of both double mutants at 7 and 21 days after cold shift compared to wild type (red circles). At 21 days after cold shift, the double mutants and *reil1-1* appeared less branched indicative of a possibly altered root branching pattern. Cultivation was in liquid MS media with 2% sucrose (w/v) adjusted to pH 5.7 (Murashige and Skoog, 1962). All photographs were taken separately as indicated by vertical white bars. In parts, black background was added to the single section of the graph for a regular and centered display of the root systems. All bars are 1 cm and indicate the slightly varying scales of single photographs.

**Supplemental Figure S4.** Differential expression of selected cold responsive genes in the roots of Col-0, and the *reil1-1 reil2-1* and *reil1-1 reil2-2* double mutants at the non-acclimated state (0 day, 20°C) and shifted to 10°C cold for 1 day or 7 days. Differential gene expression was determined relative to non-acclimated Col-0 at optimal temperature 20°C (means +/- standard error, n = 3).

The genes belonged to GO term GO:0009409, response to cold, and in part to GO term GO:0009631, cold acclimation. Note that the marker genes indicate deregulation of cold marker gene expression in non-cold acclimated *reil* double mutant roots. Analysis of variance of each of the examples indicated a significant temperature effect (*P* < 0.001, 2 way-ANOVA) and no significant interaction with the mutant effect. The mutant effect was significant (P < 0.05, 2 way-ANOVA) in the cases of *KIN2*, *COR15A*, and *COR15B*.

**Supplemental Figure S5.** Reduced transcript levels of the *reil 1* and *reil2* genes in the *reil1-1 reil2-1* and *reil1-1 reil2-2* double mutants.

Root transcript data are of this study (**Supplemental Table S2**). Leaf transcript data are from soil-grown rosette plants of *reil1-1 reil2-1* from a previous study (Supplemental Table S2 of Beine-Golovchuk et al., 2018). Log_2_-transformed transcript data were normalized to the non-acclimated Col-0 wild type, i.e. log_2_-fold changes (FC), of each transcriptome experiment, respectively. Roots (circles), leaves (diamonds).

**Supplemental Figure S6.** Functional enrichment analyses of differential gene expression in the roots of Col-0, and the *reil1-1 reil2-1* and *reil1-1 reil2-2* double mutants at the non-acclimated state (0 day, 20°C) and shifted to 10°C cold for 1 day or 7 days. Differential gene expression was determined relative to non-acclimated Col-0 at optimized temperature 20°C.

**(A)** Mean log_2_-FCs of developmental, cell wall, and root related GO terms.

**(B)** Mean log_2_-FCs of large biosynthesis related and miscellaneous GO terms.

(C = cellular component, P = biological process, F = molecular function). Significant positive or negative functional enrichments, i.e. FDR-adjusted *P* < 0.05, are indicated by asterisks, heat map color scale of log_2_-FC +1.0 (red) to -1.0 (blue). Mean log_2_-FC, z-scores, and FDR-adjusted *P*-values of gene sets from 2145 GO terms were calculated by parametric analysis of gene set enrichment (PAGE; Tian et al., 2017). The full data set is listed in Supplemental Table S3.

**Supplemental Figure S7.** Proteomic characterization of ribosome preparations obtained from root material by sucrose density gradient fractionation experiments DS1 and DS2, i.e. the root subset of Pride repository data set PXD016292. Ribosome complexes are represented by sums of label-free quantification (LFQ)-intensities of all detected RPs of 40S, 60S, and mitochondrial or plastid 30S and 50S subunits within each fraction. Non-translating immature 60S subunits are assessed by Arabidopsis homologs eIF6A and NMD3 of yeast cytosolic 60S maturation factors, means +/- standard error of the four fractionations of DS1 or DS2 (**Supplemental Table S4**).

**Supplemental Figure S8.** Distribution analysis of RPP2D **(A)**, RPP0B **(B)**, RPL3A **(C)**, and RPL3B **(D)** across the sampled ribosome fractions and analyzed conditions, DS1 orange, DS2 blue.

**Supplemental Figure S9.** Association analysis of changes in root transcript abundance and relative protein abundance in non-translating 40S and 60S (60S/80S) fractions of *reil1 reil2* mutants compared to *Arabidopsis thaliana* Col-0 wild type. DS2 (*reil1.1 reil2.1*) blue **(A, C, E)**, DS1 (*reil1.1 reil2.2*) orange **(B, D, F)**. **(A, B)** relative protein and transcript abundance determined at 20°C, i.e. at day 0, prior to cold shift; **(C, D)** relative protein abundance determined at 7 days after shift to 10°C, transcript abundance at 1 day after shift to 10°C; **(E, F)** relative protein and transcript abundance determined at 7 days after shift to 10°C.

Note that differential accumulation of cytosolic RPs and ribosome associated proteins in non-translating ribosome fractions (**Supplemental Table S6**) was matched to differential gene expression (**Supplemental Table S2** columns X to AC) according to the gene model information of the transcriptome and proteome analyses (Pride data set PXD016292). Transcript information was in part splice variant specific. The proteome analyses contained in part non-specific information due to peptides that represented multiple splice variants, and proteins of highly similar or identical protein families (**Supplemental Table S6**).

**Supplemental Table S1.** Abundance analysis of ribosome complexes (40S SSU, 60S LSU, 80S monosomes and polysomes) from roots of single-paralog *reil* mutants (*reil1-1, reil2-1, reil2-2*) and double mutants (*reil1-1 reil2-1* and *reil1-1 reil2-2*) compared to wild type Col-0 at non-acclimated (20°C, 0 day) and cold-acclimating states (10°C, 1 day, 3 days, 7 days, and 21 days).

**Supplemental Table S2.** Gene expression profiles and statistical data analyses of roots from the non-acclimated and 10°C cold acclimating *reil1-1 reil2-1* and *reil1-1 reil2-2* mutants compared to Col-0 wild type.

Genes are listed by name, description, probe identifier and probe sequence (columns A-E). The table contains differential profiles of *reil1-1 reil2-1* and *reil1-1 reil2-2* compared to Col-0 at different time points, namely at 0 day (non-acclimated state), 1 day or 7 days after cold shift (columns G-L), 2-way analyses of variance (ANOVA) (columns N-P), and pairwise heteroscedastic Student’s t-test results of differential gene expression in the mutants (column R-Y) with significance calls at *P* < 0.01 (column U and Y) and 1-way ANOVA of the cold shift response in Col-0 (column AA) with t-test results of cold acclimating Col-0 compared to non-acclimated Col-0 (column AB-AE). Differential profiles of mutants and Col-0 were compared to non-acclimated Col-0 (columns AG-AO) with respective standard errors (columns AQ-AY) and *P* values of t-tests (Column BA-BI). Note that gene expression values were averaged across redundant probes, if multiple probes per gene model were available. Only the first probe identifiers and respective probe sequences of each gene are listed. Three biological replicates (n=3) in which each replicate was a pool of roots from at least three plants grown in the same cultivation container were analyzed. Note the eight genes that were constitutively and highly expressed in both *reil* double mutants compared to wild type Col-0 regardless of the length of cold acclimation. These genes were selected by applying filters of *P* < 0.05 to each of the columns R-T and V-X. Two of these genes were also found to be consistently differentially expressed in the leaf transcriptome that was analyzed previously (Beine-Golovchuk et al., 2018). Paste values from any row below into row 3 is for visualization of differential gene expression.

**Supplemental Table S3.** Functional enrichment analysis of differential gene expression in non-acclimated (0 day) and cold-acclimating roots of *Arabidopsis thaliana* Col-0 and the *reil1-1 reil2*-*1* and *reil1-1 reil2-2* mutants at 1 day or 7 days after shift to 10°C cold. Differential gene expression was determined relative to non-acclimated Col-0 that was cultivated at 20°C, i.e. optimized cultivation temperature. Mean log_2_-fold changes (Log_2_-FC), z-scores, and *P*-values of gene set enrichments of 2145 GO terms were calculated by parametric analysis of gene set enrichment (Tian et al., 2017). GO identifiers, ontology domain, description and number of constituent genes are listed (columns A-D). Mean log_2_-FC changes are color coded by a scale ranging from +1.5 (red) to -1.5 (blue).

**Supplemental Table S4.** Proteomic characterization of ribosome preparations obtained from root material by sucrose density gradient fractionation experiments DS1 and DS2 (Selection of analyzed samples from Pride repository data set PXD016292). Ribosome complexes are characterized by sums of label-free quantification (LFQ)-intensities of all detected RPs of 40S, 60S, and organelle 30S or 50S subunits within each fraction. Non-translating immature 60S subunits are assessed by Arabidopsis homologs eIF6A and NMD3 of yeast cytosolic 60S maturation factors. Experiments DS1 and DS2 compare the ribo-proteome of 6 fractions after sucrose density gradient fractionation. Samples of *Arabidopsis thaliana* Col-0 wild type are compared to the double mutants, *reil1-1 reil2-1* (DS2) and *reil1-1 reil2-2* (DS1) at day 0 (non-acclimated) and day 7 after shift to 10°C. Sums of LFQ-intensities of cytosolic and organelle ribosome subunits are reported (* not analyzed because of low peptide count).

**Supplemental Table S5.** Fold changes (FC) of total RP abundances in the proteomic analyses of roots from the *reil1 reil2* double mutants compared to Col-0 wild type at 20°C and 7 days after shift to 10° cold. Numbers of detected proteins and sums of LFQ-abundances across all sedimentation fractions are reported.

**Supplemental Table S6.** List and specificity analysis of detected cytosolic RPs and ribosome associated proteins.

**Supplemental Table S7.** Differential accumulation of cytosolic RPs and RP associated proteins (RAPs) in non-translating ribosome fractions of the *reil1-1 reil2-1* (DS2) and *reil1-1 reil2-2* (DS1) mutants relative to *Arabidopsis thaliana* Col-0 wild type. This list contains changes that are shared between the *reil1 reil2* mutants 7 days after shift to 10°C and respective observations at 20°C prior to the shift. Supplemental Table S7A contains shared log_2_-fold changes in the non-translating 60S and 60/80S fractions after normalization of protein LFQ-abundances by LFQ-abundance sums of all detected 60S RPs. Supplemental Table S7B contains shared log_2_-fold changes in the non-translating 30/S40S fraction after normalization of protein LFQ-abundances by LFQ-abundance sums of all detected 40S RPs. Presence in mutants relative to absence in Col-0 are classified as increase, absence in mutants relative to presence in Col-0 as decrease.

## Acknowledgements

We acknowledge the research funding and facility provided by the Max Planck Society (Germany). We thank the University of Melbourne (Australia) for providing the Melbourne Research Scholarship to B.E.C. to support her PhD study. We thank Christopher Pries (MSc) for advice and support of transcriptome data preprocessing.

## Author contributions

J.K., U.R., O.B.G., W.W.H.H., and B.E.C. conceived the original research plan. B.E.C. and O.B.G performed the experiments. O.B.G. preprocessed the transcriptome data. J.K., C.B.E. and O.B.G. analyzed the preprocessed transcriptome data. M.G. and A.S. generated and preprocessed the proteome data. J.K., F.M-S., and A.A.P.F. analyzed the preprocessed proteome data. J.K. and B.E.C. wrote the manuscript with contributions of all other authors.

## References

1. Barakat, A., Szick-Miranda, K., Chang, I.F., Guyot, R., Blanc, G., Cooke, R., Delseny, M., Bailey-Serres, J. (2001). The organization of cytoplasmic ribosomal protein genes in the Arabidopsis genome. Plant Physiology 127: 398–415.

2. Basu, U., Si, K., Warner, J.R., Maitra, U. (2001). The Saccharomyces cerevisiae TIF6 gene encoding translation initiation factor 6 is required for 60S ribosomal subunit biogenesis. Mol Cell Biol 21: 1453–1462.

3. Beine-Golovchuk, O., Firmino, A.A.P., Dabroska, A., Schmidt, S., Erban, A., Walter, D., Zuther, E., Hincha, D.K., Kopka, J. (2018). Plant temperature acclimation and growth rely on cytosolic ribosome factor homologs. Plant Physiology 176: 2251–2276.

4. Benjamini, Y., Hochberg, Y. (1995). Controlling the false discovery rate: a practical and powerful approach to multiple testing. Journal of the Royal Statistical Society (Series B) 57:289–300.

5. Boyes, D.C., Zayed, A.M., Ascenzi, R., McCaskill, A.J., Hoffman, N.E., Davis, K.R., Gorlach, J. (2001). Growth stage-based phenotypic analysis of Arabidopsis: a model for high throughput functional genomics in plants. Plant Cell 13: 1499–1510.

6. Brady, S.M., Orlando, D.A., Lee, J-Y., Wang, J.Y., Koch, J., Dinneny, J.R., Mace, D. et al. (2007). A high-resolution root spatiotemporal map reveals dominant expression patterns. Science 318: 801–806.

7. Browning, K.S., Bailey-Serres, J. (2015). Mechanism of cytoplasmic mRNA translation. Arabidopsis Book 13: e0176

8. Browning, K.S., Gallie, D.R., Hershey, J.W., Hinnebusch, A.G., Maitra, U., Merrick, W.C., Norbury, C. (2001). Unified nomenclature for the subunits of eukaryotic initiation factor 3. Trends Biochem Sci 26: 284.

9. Burks, E.A., Bezerra, P.P., Le, H., Gallie, D.R., Browning, K.S. (2001). Plant initiation factor 3 subunit composition resembles mammalian initiation factor 3 and has a novel subunit. Journal of Biological Chemistry 276: 2122–2131.

10. Cheng, Z., Mugler, C.F., Keskin, A., Hodapp, S., Chan, L.Y.L., Weis, K., Mertins, P., Regev, A., Jovanovic, M., Brar, G.A. (2019). Small and large ribosomal subunit deficiencies lead to distinct gene expression signatures that reflect cellular growth rate. Molecular Cell 73: 36–47.

11. Cook, D., Fowler, S., Fiehn, O., Thomashow, M.F. (2004). A prominent role for the CBF cold response pathway in configuring the low-temperature metabolome of *Arabidopsis*. Proc Natl Acad Sci USA 101: 15243–15248.

12. Dong Y., Teleman, A.A., Jedmowski, C., Wirtz, M., Hell, R. (2019). The Arabidopsis THADA homologue modulates TOR activity and cold acclimation. Plant Biology 21: 77–83.

13. Du, Z., Zhou, X., Ling, Y., Zhang, Z., Su, Z. (2010). AgriGO: a GO analysis toolkit for the agricultural community. Nucleic Acids Res 38: W64–W70.

14. Durut, N., Abou-Ellail M., Pontvianne, F., Das, S., Kojima, H., Ukai, S., de Bures, P. et al. (2014). A duplicated NUCLEOLIN gene with antagonistic activity is required for chromatin organization of silent 45D rDNA in Arabidopsis. Plant Cell 26: 1330–1344.

15. Erde, J., Loo, R. R. O., Loo, J. A. (2014). Enhanced FASP (eFASP) to increase proteome coverage and sample recovery for quantitative proteomic experiments. J Proteome Res 13: 1885–1895.

16. Gamalinda, M., Jakovljevic, J., Babiano, R., Talkish, J., de la Cruz, J., Woolford, J.L. (2013). Yeast polypeptide exit tunnel ribosomal proteins L17, L35 and L37 are necessary to recruit late-assembling factors required for 27SB pre-rRNA processing. Nucleic Acids Res 41:1965–1983.

17. Genuth, N. R., Barna, M. (2018). Heterogeneity and specialized functions of translation machinery: from genes to organisms. Nature Reviews Genetics 19: 431–452.

18. Gnanasundram, S.V., Kos-Braun, I.C., Kǒs, M. (2019). At least two molecules of the RNA helicase Has1 are simultaneously present in pre-ribosomes during ribosome biogenesis. Nucleic Acids Res. 47: 10852–10864.

19. Greber, B.J., Boehringer, D., Montellese, C., Ban, N. (2012). Cryo-EM structures of Arx1 and maturation factors Rei1 and Jjj1 bound to the 60S ribosomal subunit. Nature Struct. Mol. Biol. 19: 1228–1234

20. Greber, B.J., Gethardy, S., Leitner, A., et al. (2016). Insertion of the biogenesis factor Rei1 probes the ribosomal tunnel during 60S maturation. Cell 164: 91–102.

21. Harnpicharnchai, P., Jakovljevic, J., Horsey, E., Miles, T., Roman, J., Rout, M., Meagher, D., Imai, B., Guo, Y., Brame, C.J., Shabanowitz, J., Hunt, D.F., Woolford, J.L. Jr. (2001) Composition and functional characterization of yeast 66S ribosome assembly intermediates. Molecular Cell 8: 505–515.

22. Ho, J.H., Kallstrom, G., Johnson, A.W. (2000). Nmd3p is a Crm1p-dependent adapter protein for nuclear export of the large ribosomal subunit. J Cell Biol 151:1057–1066.

23. Holt, C.E., Schuman, E.M. (2013). The central dogma decentralized: new perspectives on RNA function and local translation in neurons. Neuron 80: 648–657.

24. Hummel, M., Dobrenel, T., Cordewener, J. J. H. G., Davanture, M., Meyer, C., Smeekens, S. J. C. M., Bailey-Serres, J., America, T.A., Hanson, J. (2015). Proteomic LC-MS analysis of Arabidopsis cytosolic ribosomes: Identification of ribosomal protein paralogs and re-annotation of the ribosomal protein genes. J. Proteomics 128: 436–449.

25. Iwase, M., Toh-e, A. (2004) Ybr267w is a new cytoplasmic protein belonging to the mitotic signaling network of Saccharomyces cerevisiae. Cell Structure and Function 29: 1–15.

26. Jung, J.K.H., McCouch, S. (2013). Getting to the roots of it: genetic and hormonal control of root architecture. Front Plant Sci 4: 186

27. Kaplan, F., Kopka, J., Haskell, D.W., Zhao, W., Schiller, K.C., Gatzke, N., Sung, D.Y., Guy, C.L. (2004). Exploring the temperature-stress metabolome of *Arabidopsis*. Plant Physiology 136: 4159–4168.

28. Kaplan, F., Kopka, J., Sung, D.Y., Zhao, W., Popp, M., Porat, R., Guy, C.L. (2007). Transcript and metabolite profiling during cold acclimation of Arabidopsis reveals an intricate relationship of cold-regulated gene expression with modifications in metabolite content. The Plant Journal 50: 967–981.

29. Kater, L., Thoms, M., Barrio-Garcia, C., Cheng, J., Ismail, S., Ahmed, Y. L., Bange, G., Kressler, D., Berninghausen, O., Sinning, I., Hurt, E., Beckmann, R. (2017). Visualizing the assembly pathway of nucleolar pre-60S ribosomes. Cell, 171: 1599–1610.

30. Kim, Y., Zhang, H., Scholl, R.L. (1990). Two evolutionarily divergent genes encode a cytoplasmic ribosomal protein of *Arabidopsis thaliana*. Gene 93: 177–182.

31. Krokowski, D., Tchórzewski, M., Boguszewska, A., Grankowski, N. (2005). Acquisition of a stable structure by yeast ribosomal P0 protein requires binding of P1A-P2B complex: in vitro formation of the stalk structure. Biochim. Biophys. Acta 1724: 59–70.

32. Krokowski, D., Boguszewska, A., Abramczyk, D., Liljas, A., Tchórzewski, M., Grankowski, N. (2006). Yeast ribosomal P0 protein has two separate binding sites for P1/P2 proteins. Mol. Microbiol. 60(2): 386–400.

33. Krokowski, D., Gaccioli, F., Majumder, M., Mullins, M.R., Yuan, C.L., Papadopoulou, B., Merrick, W.C., Komar, A.A., Taylor, D., Hatzoglou, M. (2011). Characterization of hibernating ribosomes in mammalian cells. Cell Cycle 10: 2691–2702.

34. Lebreton, A., Saveanu, C., Decourty, L., Rain, J-C., Jacquier, A., Fromont-Racine, M. (2006). A functional network involved in the recycling of nucleocytoplasmic pre-60S factors. Journal of Cell Biology 173: 349–360.

35. Lee, J-Y., Colinas, J., Wang, J.Y., Mace, D., Ohler, U., Benfey, P.N. (2006). Transcriptional and posttranscriptional regulation of transcription factor expression in Arabidopsis roots. PNAS 103: 6055–6060.

36. Li, L., Nelson, C. J., Trösch, J., Castleden, I., Huang, S., & Millar, A. H. (2017). Protein degradation rate in Arabidopsis thaliana leaf growth and development. The Plant Cell 29: 207–228.

37. Liljas, A, Sanyal, S (2018). The enigmatic ribosomal stalk. Quarterly Reviews of Biophysics 51, e12, 1–10.

38. Lo, K.Y., Li, Z.H., Bussiere, C., Bresson, S., Marcotte, E.M., Johnson, A.W. (2010) Defining the pathway of cytoplasmic maturation of the 60S ribosomal subunit. Molecular Cell 39: 196–208.

39. Ma, C., Wu, S., Li, N., Chen, Y., Yan, K., Li, Z., Zheng, L., Lei, J., Woolford, J.L. Jr., Gao, N. (2017). Structural snapshot of cytoplasmic pre-60S ribosomal particles bound by Nmd3, Lsg1, Tif6 and Reh1. Nat Struct Mol Biol 24: 214–220.

40. Maruyama, K., Sakuma, Y., Kasuga, M., Ito, Y., Seki, M., Goda, H., Shimada, Y., Yoshida, S., Shinozaki, K., Yamaguchi-Shinozaki, K. (2004). Identification of cold-inducible downstream genes of the Arabidopsis DREB1A/CBF3 transcriptional factor using two microarray systems. The Plant Journal 38: 982–993.

41. Martin, T.E., Hartwell, L.H. (1970). Resistance of active yeast ribosomes to dissociation by KCl. Journal of Biological Chemistry 245: 1504–1506.

42. Matsuo, Y., Granneman, S., Thoms, M., Manikas, R.G., Tollervey, D., Hurt, E. (2014). Coupled GTPase and remodelling ATPase activities form a checkpoint for ribosome export. Nature 505:112–116.

43. Meinke, D.W. (2019). Genome-wide identification of EMBRYO-DEFECTIVE (EMB) genes required for growth and development in Arabidopsis. New Phytologist doi: 10.1111/nph.16071.

44. Mitroshin, I. V., Garber, M. B., Gabdulkhakov, A. G. (2016) Investigation of Structure of the Ribosomal L12/P Stalk. Biochemistry (Moscow), 81(13): 1589–1601.

45. Morita, D., Miyoshi, K., Matsui, Y., Toh-e, A., Shinkawa, H., Miyakawa, T., Mizuta, K. (2002). Rpf2p, an evolutionarily conserved protein, interacts with ribosomal protein L11 and is essential for the processing of 27 SB Pre-rRNA to 25 S rRNA and the 60 S ribosomal subunit assembly in *Saccharomyces cerevisiae*. J Biol Chem 277: 28780–28786.

46. Pagter, M., Alpers, J., Erban, A., Kopka, J., Zuther, E., Hincha, D.K. (2017). Rapid transcriptional and metabolic regulation of the deacclimation process in cold acclimated *Arabidopsis thaliana*. BMC Genomics 18: 731

47. Parnell, K.M., Bass, B.L. (2009). Functional redundancy of yeast proteins Reh1 and Rei1 in cytoplasmic 60S subunit maturation. Mol Cell Biol 29: 4014–4023.

48. Perez-Riverol, Y., Csordas, A., Bai, J., Bernal-Llinares, M., Hewapathirana, S., Kundu, D. J., Inuganti, A., Griss, J., Mayer, G., Eisenacher, M., Pérez, E., Uszkoreit, J., Pfeuffer, J., Sachsenberg, T., Yılmaz, Ş., Tiwary, S., Cox, J., Audain, E., Walzer, M., Jarnuczak, A.F., Ternent, T., Brazma, A., Vizcaíno, J.A. (2019). The PRIDE database and related tools and resources in 2019: Improving support for quantification data. Nucleic Acids Res. 47: D442–D450.

49. Rosso, M.G., Li, Y., Strizhov, N., Reiss, B., Dekker, K., Weisshaar, B. (2003). An Arabidopsis thaliana T-DNA mutagenized population (GABI-Kat) for flanking sequence tag-based reverse genetics. Plant Molecular Biology 53: 247–259.

50. Rugen, N., Straube, H., Franken, L. E., Braun, H. P., Eubel, H. (2019). Complexome profiling reveals association of PPR proteins with ribosomes in the mitochondria of plants. Mol. Cell. Proteomics 18: 1345–1362.

51. Sáez-Vásquez, J., Delseny, M. (2019). Ribosome biogenesis in plants: from functional 45S ribosomal DNA organization to ribosome assembly factors. Plant Cell 31: 1945–1967.

52. Schmidt, S., Dethloff, F., Beine-Golovchuk, O., Kopka, J. (2013). The REIL1 and REIL2 proteins of *Arabidopsis thaliana* are required for leaf growth in the cold. Plant Physiology 163: 1623–1639.

53. Schmidt, S., Dethloff, F., Beine-Golovchuk, O., Kopka, J. (2014). REIL proteins of *Arabidopsis thaliana* interact in yeast-2-hybrid assays with homologs of the yeast Rlp24, Rpl24A, Rlp24B, Arx1, and Jjj1 proteins. Plant Signal Behaviour 9: e28224.

54. Scholl, R.I., Ma, S.T., Ware, D.H. (2000). Seed and molecular resources for Arabidopsis. Plant Physiology 124: 1477–1480.

55. Salih, K.J., Duncan, O., Li, L., Troesch, J., Millar, A.H. (2019). Refining the composition of the *Arabidopsis thaliana* 80S cytosolic ribosome. bioRxiv 764316

56. Shi, Z., Fujii, K., Kovary, K. M., Genuth, N. R., Röst, H. L., Teruel, M. N., Barna, M. (2017). Heterogeneous ribosomes preferentially translate distinct subpools of mRNAs genome-wide. Molecular Cell 67: 71–83.

57. Simsek, D., Tiu, G. C., Flynn, R. A., Byeon, G. W., Leppek, K., Xu, A. F., Chang, H.Y., Barna, M. (2017). The mammalian ribo-interactome reveals ribosome functional diversity and heterogeneity. Cell 169: 1051–1065.

58. Sormani, R., Masclaux-Daubresse, C., Daniel-Vedele, F., Chardon, F. (2011) Transcriptional regulation of ribosome components are determined by stress according to cellular compartments in *Arabidopsis thaliana*. PLoS ONE 6: e28070

59. Stitt, M., Hurry, V. (2002). A plant for all seasons: alterations in photosynthetic carbon metabolism during cold acclimation in Arabidopsis. Current Opinion in Plant Biology 5: 199–206

60. Swart, C., Martínez-Jaime, S., Gorka, M., Zander, K., Graf, A. (2018). Hit-Gel: Streamlining in-gel protein digestion for high-throughput proteomics experiments. Sci Rep. 8: 8582.

61. The UniProt Consortium (2017). UniProt: the universal protein knowledgebase. Nucleic Acids Res. 45: D158–D169.

62. Uesono, Y., Toh-e, A. (2002). Transient inhibition of translation initiation by osmotic stress. Journal of Biological Chemistry 277: 13845–13855.

63. Van den Elzen, A.M.G., Schuller A., Green, R., Seraphin, B. (2014). Dom34-Hbs1 mediated dissociation of inactive 80S ribosomes promotes restart of translation after stress. The EMBO Journal 33: 265–276.

64. Waltz, F., Nguyen, T. T., Arrivé, M., Bochler, A., Chicher, J., Hammann, P., Kuhn L., Quadrado M., Mireau H., Hashem, Y., Giegé, P. (2019). Small is big in Arabidopsis mitochondrial ribosome. Nature Plants 5: 106–117

65. Weis, B.L., Kovacevic, J., Missbach, S., Schleiff, E. (2015). Plant-specific features of ribosome biogenesis. Trends in Plant Science 20: 729–740.

66. Tian, T., Liu, Y., Yan, H., You, Q., Yi, X., Du, Z., Xu, W., Su, Z. (2017). AgriGO v2.0: a GO analysis toolkit for the agricultural community, 2017 update. Nucleic Acids Research 45: W122–W129.

67. Van Buskirk, H.A., Thomashow, M.F. (2006). *Arabidopsis* transcription factors regulating cold acclimation. Physiologia Plantarum 126: 72–80.

68. Xiong, Y., McCormack, M., Li, L., Hall, Q., Xiang, C., Sheen, J. (2013) Glucose-TOR signalling reprograms the transcriptome and activates meristems. Nature 496: 181–186.

69. Xue, S., Barna, M. (2012). Specialized ribosomes: A new frontier in gene regulation and organismal biology. Nature Reviews Molecular Cell Biology, 13: 355–369.

70. Yu, H., Kong, X., Huang, H., Wu, W., Park, J., Yun, D.J., Lee, B.H., Shi, H., Zhu, J.K. (2020). STCH4/REIL2 confers cold stress tolerance in Arabidopsis by promoting rRNA processing and CBF protein translation. Cell Reports 30: 229–242.

71. Zhang, W., Zhang, J., Xu, C., Li, N., Liu, H., Ma, J., Zhu, Y., Xie, H. (2012). LFQuant: a label-free fast quantitative analysis tool for high-resolution LC-MS/MS proteomics data. Proteomics 12: 3475–3484.

72. Zylber, E.A., Penman, S. (1970). The effect of high ionic strength on monomers, polyribosomes, and puromycin-treated polyribosomes. Biochim Biophys Acta 204: 221–229.

