## Supplemental Figures S1- S8 for "Arabidopsis REI-LIKE proteins activate ribosome biogenesis during cold acclimation"

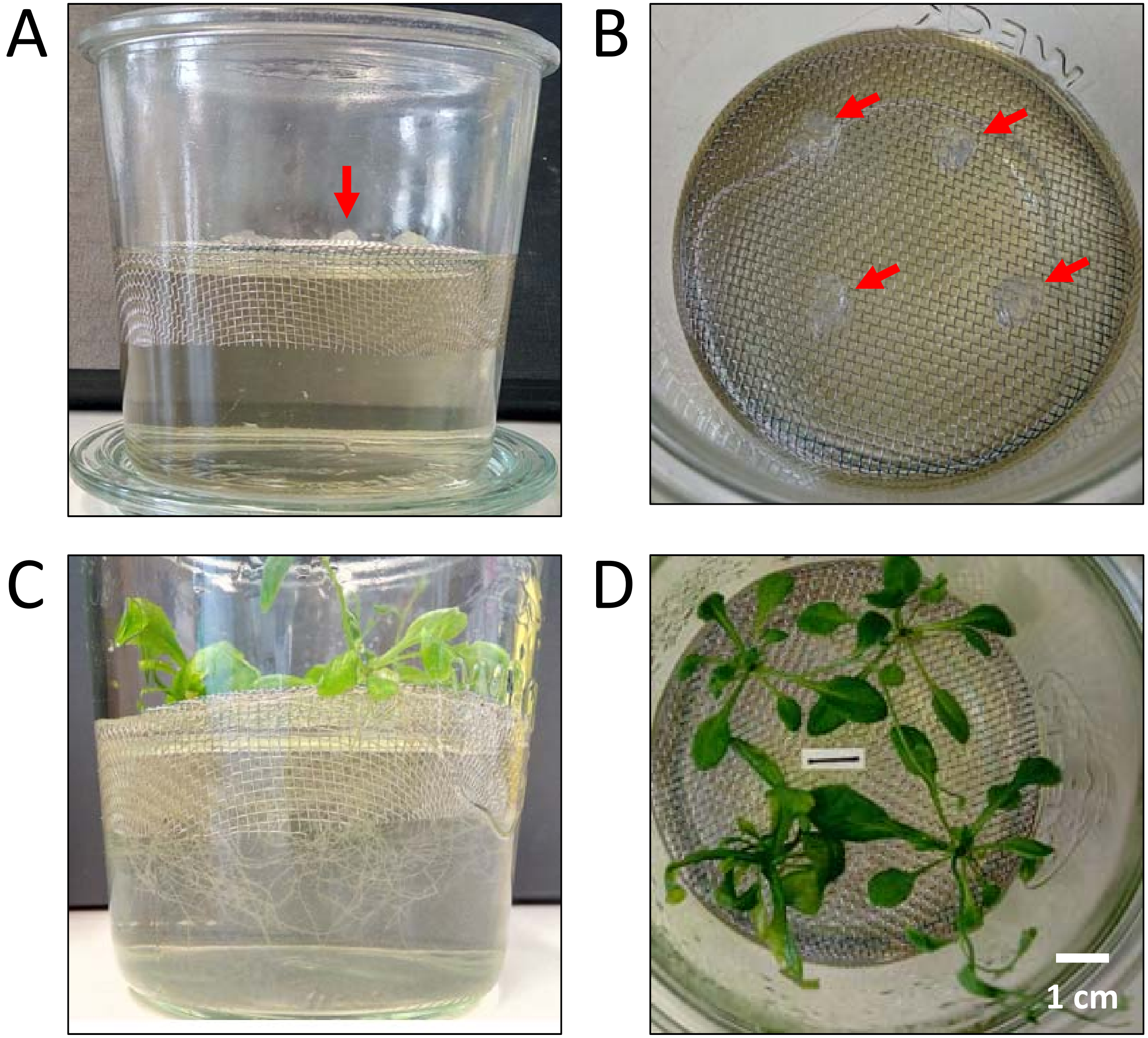
A
B
C
D
1 cm

**Supplemental Figure S1.** Hydroponic growth system of *Arabidopsis thaliana* Col-0 wild type and mutants. The system was feasible for temperature shift experiments in the vegetative phase and for rapid sampling of pooled material of whole root or shoot systems.

**A)** Assembled growth system with stainless steel mesh adjusted level with the surface of ~ 250 mL liquid Murashige Skoog-medium with 2% sucrose. One of the four agar blocks for seed germination is indicated (red arrow, view from the side).

**B)** Assembled growth system with stainless steel mesh adjusted level with the surface of the liquid. The four agar blocks for seed germination are indicated (red arrows, view from the top). Each block contains a single seed.

**C)** Assembled growth system with *Arabidopsis thaliana* Col-0 wild type plants at approximate ten-leaf stage after 4 weeks in 16 h/ 8 h long day conditions at 20 °C during the day and 18 °C during night (view from the side).

**D)** Assembled growth system with *Arabidopsis thaliana* Col-0 wild type plants at approximately ten-leaf stage (view from the top).

The clamped-on glass lids were opened and steril conditions compromised for the photographs.

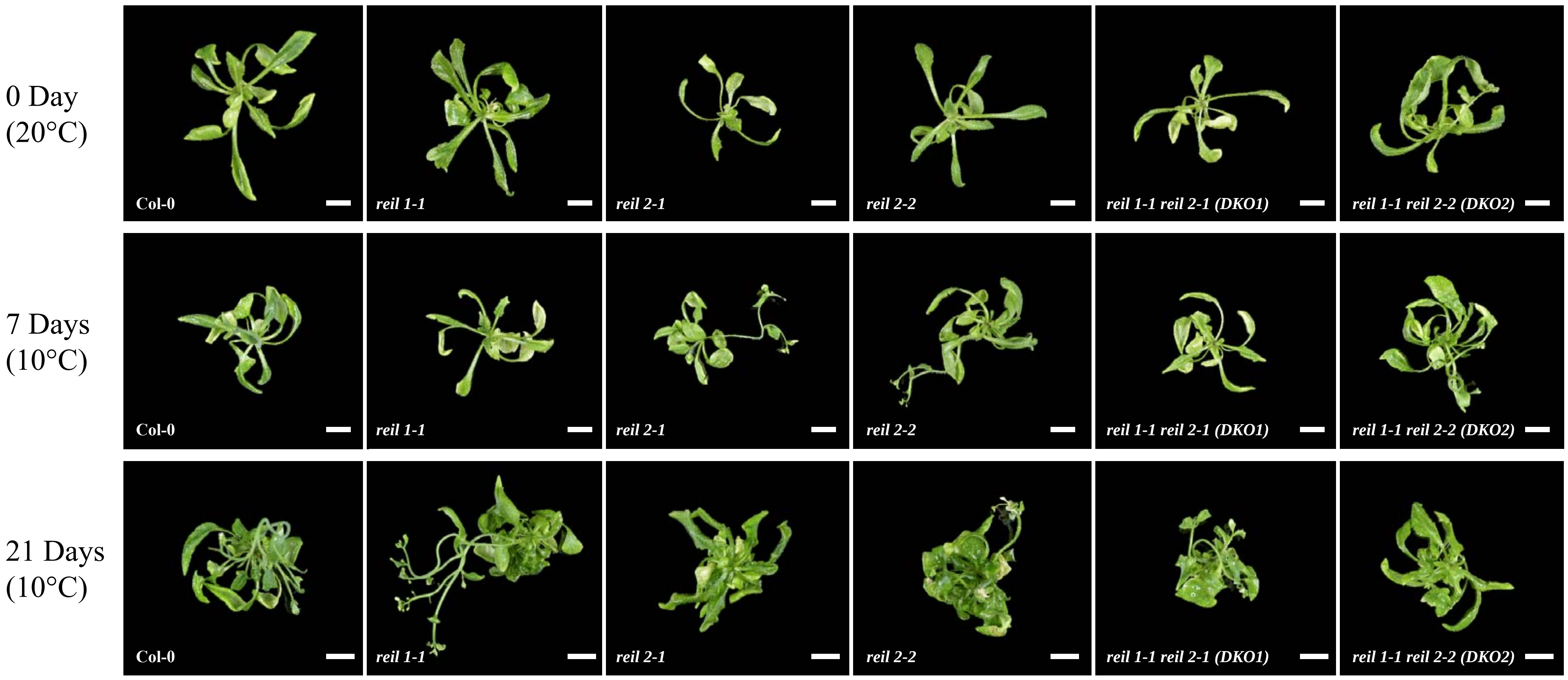
0 Day
(20°C)
7 Days
(10°C)
21 Days
(10°C)
Col-0
reil 1-1
reil 2-1
reil 2-2
reil 1-1 reil 2-1 (DKO1)
reil 1-1 reil 2-2 (DKO2)

**Supplemental Figure S2.** Exemplary documentation of shoot systems of hydroponically cultivated *Arabidopsis thaliana* wild type (Col-0), of the single mutants, *reil1-1, reil2-1, reil2-2,* and of the double mutants, *reil1-1 reil2-1 (DKO1)* and *reil1-1 reil2-2 (DKO2),* before (0 day) and 7 or 21 days after shift from 20°C (day)/ 18°C (night) to 10°C (day) and 8°C (night).
Note that at 21 days after cold shift shoots and inflorescences of *reil2-1, reil2-2,* and the double mutants, *reil1-1 reil2-1 (DKO1)* and *reil1-1 reil2-2 (DKO2),* are smaller than wild type and the *reil1-1* mutant. Growth of the double mutants is not arrested under the conditions of this hydroponic system. Cultivation was in liquid MS media with 2% sucrose (w/v) adjusted to pH 5.7 (Murashige and Skoog, 1962). All photographs were taken separately as indicated by vertical white bars. In parts, black background was added to the single sections of the graph for a regular and centered display of the shoot systems. All bars are 1 cm and indicate the slightly varying scales of single photographs.

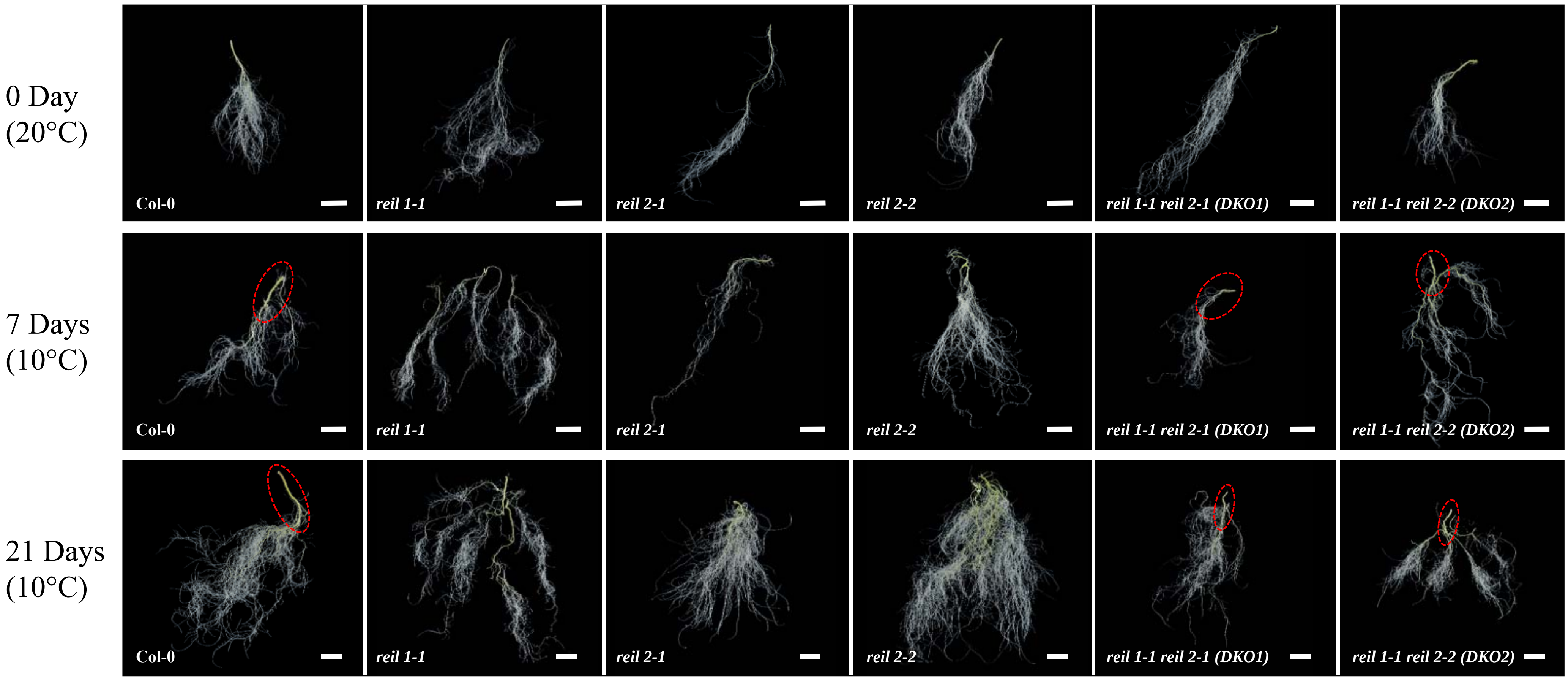
0 Day
(20°C)
7 Days
(10°C)
21 Days
(10°C)
Col-0
reil 1-1
reil 2-1
reil 2-2
reil 1-1 reil 2-1 (DKO1)
reil 1-1 reil 2-2 (DKO2)

**Supplemental Figure S3.** Exemplary documentation of hydroponic root systems of *Arabidopsis thaliana* wild type (Col-0), of the single mutants, *reil1-1*, *reil2-1*, *reil2-2*, and of the double mutants, *reil1-1 reil2-1* (*DKO1*) and *reil1-1 reil2-2* (*DKO2*), before (0 day) and 7 or 21 days after shift from 20°C (day)/ 18°C (night) to 10°C (day) and 8°C (night).
Roots were cut at the hypocotyl to root transition. Root systems of single plants were carefully prepared with minimal wounding from joined cultivations of four plants in single containers. Preparation of complete root systems from single plants was not possible. Photographs may therefore show in parts incomplete root systems. Note the shortened primary root of both double mutants at 7 and 21 days after cold shift compared to wild type. The respective primary root sections are indicated by red circles. At 21 days after cold shift, the double mutants and *reil1-1* appeared less branched indicative of an possibly altered root branching pattern. Cultivation was in liquid MS media with 2% sucrose (w/v) adjusted to pH 5.7 (Murashige and Skoog, 1962). All photographs were taken separately as indicated by vertical white bars. In parts, black background was added to the single section of the graph for a regular and centered display of the root systems. All bars are 1 cm and indicate the slightly varying scales of single photographs.

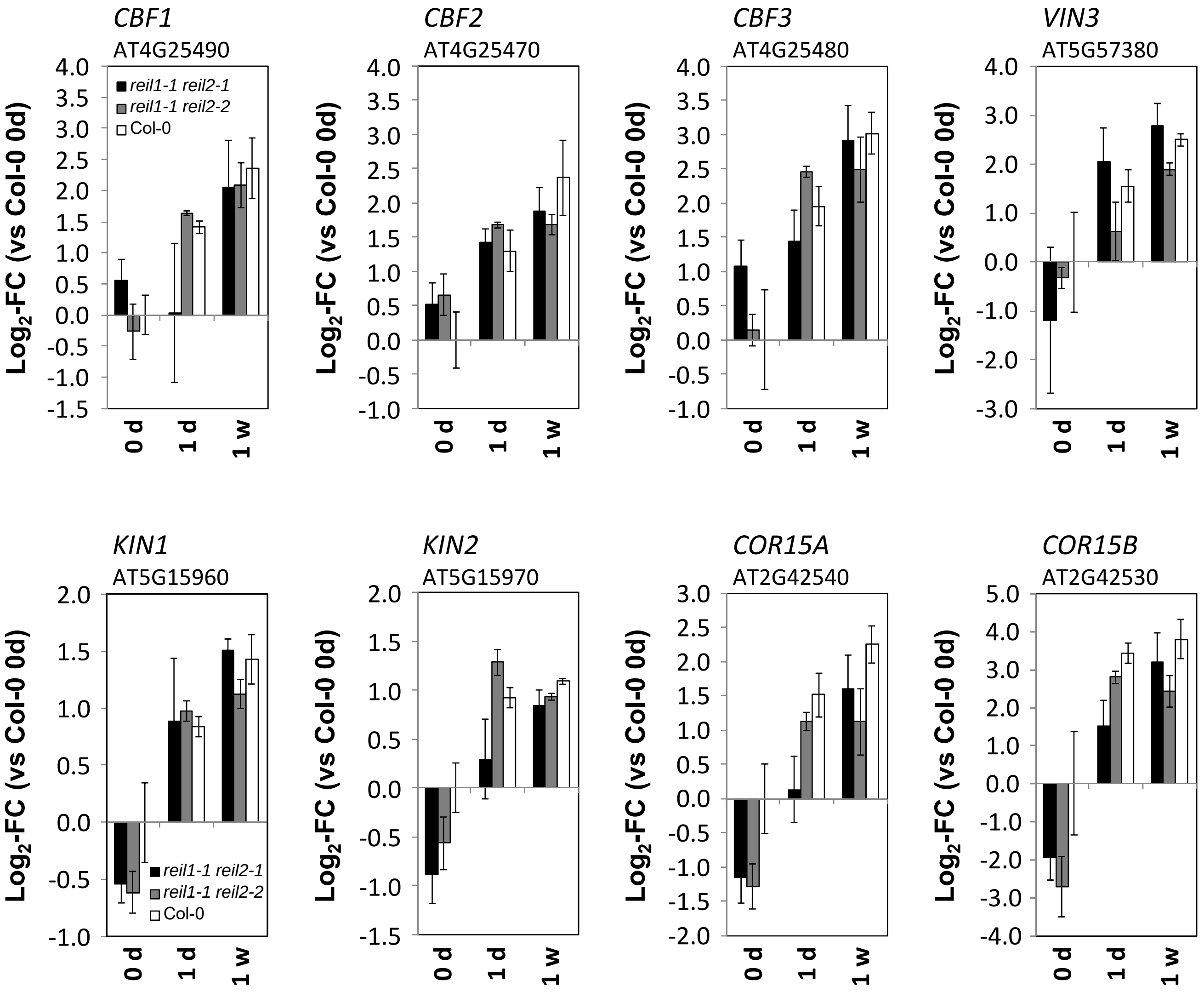

CBF1
AT4G25490
CBF2
AT4G25470
CBF3
AT4G25480
VIN3
AT5G57380
KIN1
AT5G15960
KIN2
AT5G15970
COR15A
AT2G42540
COR15B
AT2G42530
reil1-1 reil2-1
reil1-1 reil2-2
Col-0
Log2-FC (vs Col-0 0d)
0 d
1 d
1 w

**Supplemental Figure S4.**

Differential expression of selected cold responsive genes in the roots of Col-0, and the *reil1-1 reil2-1* and *reil1-1 reil2-2* double mutants at the non-acclimated state (0 day, 20°C) and shifted to 10°C cold for 1 day or 1 week. Differential gene expression was determined relative to non-acclimated Col-0 at optimal temperature 20°C.

The genes belonged to GO term GO:0009409, response to cold, and in part to GO term GO:0009631, cold acclimation. Note that the marker genes indicate deregulation of cold marker gene expression in non-cold acclimated *reil* double mutant roots. Analysis of variance of each of the examples indicated a significant temperature effect ($P < 0.001$, 2 way-ANOVA) and no significant interaction with the mutant effect. The mutant effect was significant ($P < 0.05$, 2 way-ANOVA) in the cases of *KIN2*, *COR15A*, and *COR15B*.

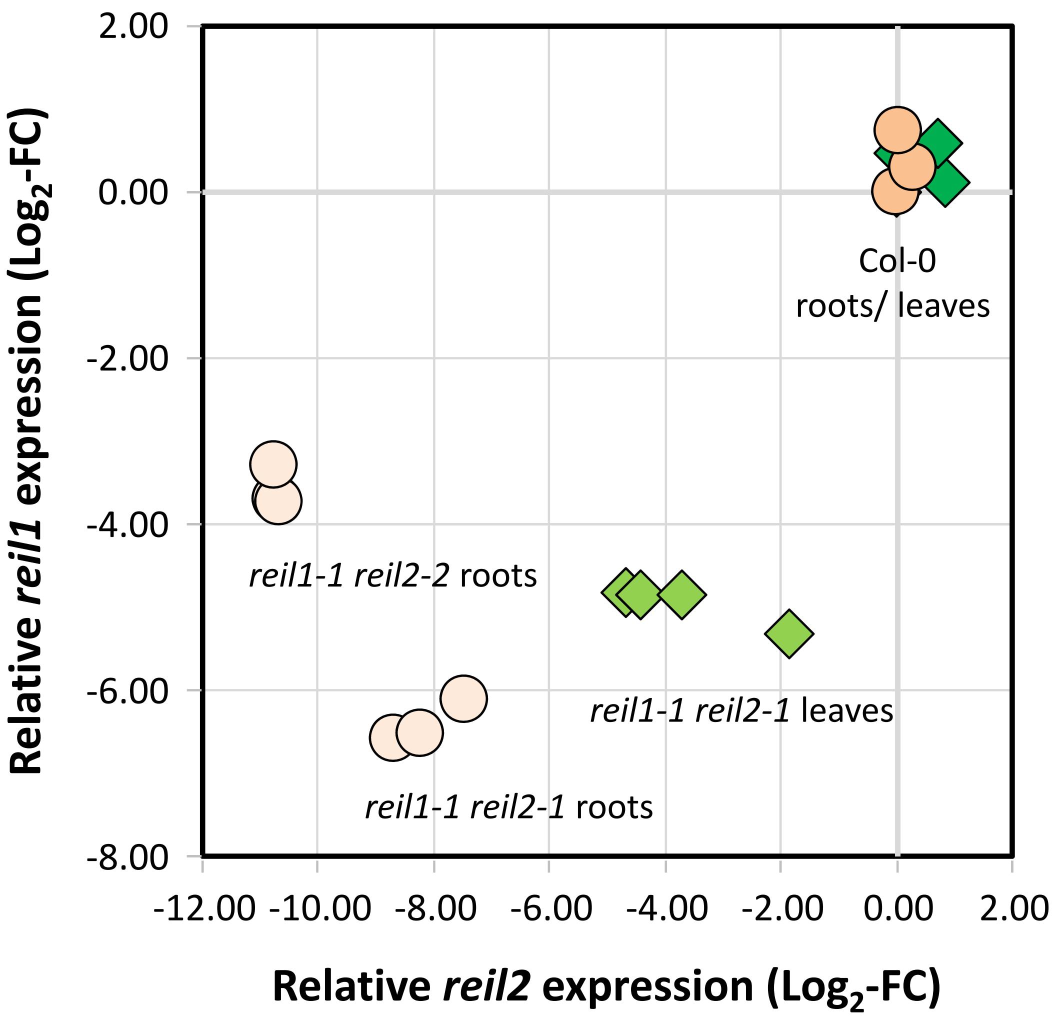

2.00
0.00
-2.00
-4.00
-6.00
-8.00
-12.00
-10.00
-8.00
-6.00
-4.00
-2.00
0.00
2.00
Relative *reil1* expression (Log2-FC)
Relative *reil2* expression (Log2-FC)
Col-0
roots/ leaves
*reil1-1 reil2-2* roots
*reil1-1 reil2-1* leaves
*reil1-1 reil2-1* roots

**Supplemental Figure S5.** Reduced transcript levels of the *reil 1* and *reil2* genes in the *reil1-1 reil2-1* and *reil1-1 reil2-2* double mutants.
Root transcript data are of this study (**Supplemental Table S2**). Leaf transcript data are from soil-grown rosette plants of *reil1-1 reil2-1* from a previous study (Supplemental Table S2 of Beine-Golovchuk et al., 2018). $Log_2$-transformed transcript data were normalized to the non-acclimated Col-0 wild type, i.e. $log_2$-fold changes (FC), of each transcriptome experiment, respectively. Roots (circles), leaves (diamonds).

**A**

| GO | (Onthology) | Description | Genes | Col-0 0 d | Col-0 1 d | Col-0 1 w | *reil1-1 reil2-1* 0 d | *reil1-1 reil2-1* 1 d | *reil1-1 reil2-1* 1 w | *reil1-1 reil2-2* 0 d | *reil1-1 reil2-2* 1 d | *reil1-1 reil2-2* 1 w |
|---|---|---|---|---|---|---|---|---|---|---|---|---|
| GO:0048589 | P | developmental growth | 211 | | -0.08 * | -0.21 * | -0.18 * | -0.13 * | -0.20 * | -0.11 * | -0.12 * | -0.11 * |
| GO:0048468 | P | cell development | 158 | | -0.06 | -0.22 * | -0.15 * | -0.16 * | -0.24 * | -0.12 * | -0.16 * | -0.13 * |
| GO:0021700 | P | developmental maturation | 47 | | -0.20 * | -0.40 * | -0.23 * | -0.29 * | -0.38 * | -0.28 * | -0.36 * | -0.26 * |
| GO:0048469 | P | cell maturation | 36 | | -0.23 * | -0.47 * | -0.30 * | -0.31 * | -0.45 * | -0.35 * | -0.39 * | -0.30 * |
| GO:0010015 | P | root morphogenesis | 91 | | -0.11 | -0.28 * | -0.17 * | -0.18 * | -0.28 * | -0.17 * | -0.20 * | -0.15 * |
| GO:0010053 | P | root epidermal cell differentiation | 49 | | -0.21 * | -0.45 * | -0.29 * | -0.33 * | -0.45 * | -0.33 * | -0.38 * | -0.26 * |
| GO:0048765 | P | root hair cell differentiation | 36 | | -0.23 * | -0.47 * | -0.30 * | -0.31 * | -0.45 * | -0.35 * | -0.39 * | -0.30 * |
| GO:0010054 | P | trichoblast differentiation | 43 | | -0.19 * | -0.45 * | -0.29 * | -0.30 * | -0.45 * | -0.35 * | -0.37 * | -0.25 * |
| GO:0048764 | P | trichoblast maturation | 36 | | -0.23 * | -0.47 * | -0.30 * | -0.31 * | -0.45 * | -0.35 * | -0.39 * | -0.30 * |
| GO:0071554 | P | cell wall organization or biogenesis | 233 | | -0.22 * | -0.30 * | -0.20 * | -0.24 * | -0.29 * | -0.13 * | -0.10 * | -0.15 * |
| GO:0016757 | F | transferase activity, transferring glycosyl | 484 | | -0.10 * | -0.12 * | -0.12 * | -0.10 * | -0.10 * | -0.08 * | -0.08 * | -0.09 * |
| GO:0016758 | F | transferase activity, transferring hexosyl | 303 | | -0.13 * | -0.13 * | -0.13 * | -0.12 * | -0.12 * | -0.11 * | -0.11 * | -0.13 * |

**B**

| GO | (Onthology) | Description | Genes | Col-0 0 d | Col-0 1 d | Col-0 1 w | *reil1-1 reil2-1* 0 d | *reil1-1 reil2-1* 1 d | *reil1-1 reil2-1* 1 w | *reil1-1 reil2-2* 0 d | *reil1-1 reil2-2* 1 d | *reil1-1 reil2-2* 1 w |
|---|---|---|---|---|---|---|---|---|---|---|---|---|
| GO:0009058 | P | biosynthetic process | 3716 | | 0.04 * | 0.01 * | -0.05 * | 0.05 * | 0.02 * | 0.01 * | 0.03 * | 0.01 * |
| GO:0009059 | P | macromolecule biosynthetic process | 2505 | | 0.06 * | 0.03 * | -0.04 * | 0.07 * | 0.04 * | 0.02 * | 0.05 * | 0.03 * |
| GO:0034645 | P | cellular macromolecule biosynthetic pro | 2483 | | 0.07 * | 0.03 * | -0.04 * | 0.07 * | 0.04 * | 0.02 * | 0.05 * | 0.04 * |
| GO:0044249 | P | cellular biosynthetic process | 3536 | | 0.05 * | 0.02 * | -0.05 * | 0.06 * | 0.02 * | 0.01 * | 0.04 * | 0.02 * |
| GO:0016137 | P | glycoside metabolic process | 86 | | 0.01 | -0.04 | 0.17 * | 0.25 * | 0.17 * | 0.24 * | 0.21 * | 0.13 * |
| GO:0030145 | F | manganese ion binding | 30 | | 0.04 | 0.00 | -0.56 * | 0.29 * | 0.27 * | -0.41 * | 0.47 * | 0.22 * |

**Supplemental Figure S6.** Functional enrichment analyses of differential gene expression in the roots of Col-0, and the *reil1-1 reil2-1* and *reil1-1 reil2-2* double mutants at the non-acclimated state (0 day, 20°C) and shifted to 10°C cold for 1 day or 1 week. Differential gene expression was determined relative to non-acclimated Col-0 at optimized temperature 20°C.

**(A)** Mean $log_2$-FCs of developmental, cell wall, and root related GO terms.

**(B)** Mean $log_2$-FCs of large biosynthesis related and miscellaneous GO terms.

(C = cellular component, P = biological process, F = molecular function). Significant positive or negative functional enrichments, i.e. FDR-adjusted $P < 0.05$, are indicated by asterisks, heat map color scale of $log_2$-FC +1.0 (red) to -1.0 (blue). Mean $log_2$-FC, z-scores, and FDR-adjusted *P*-values of gene sets from 2145 GO terms were calculated by parametric analysis of gene set enrichment (PAGE; Tian et al., 2017). The full data is listed in Supplemental Table S3.

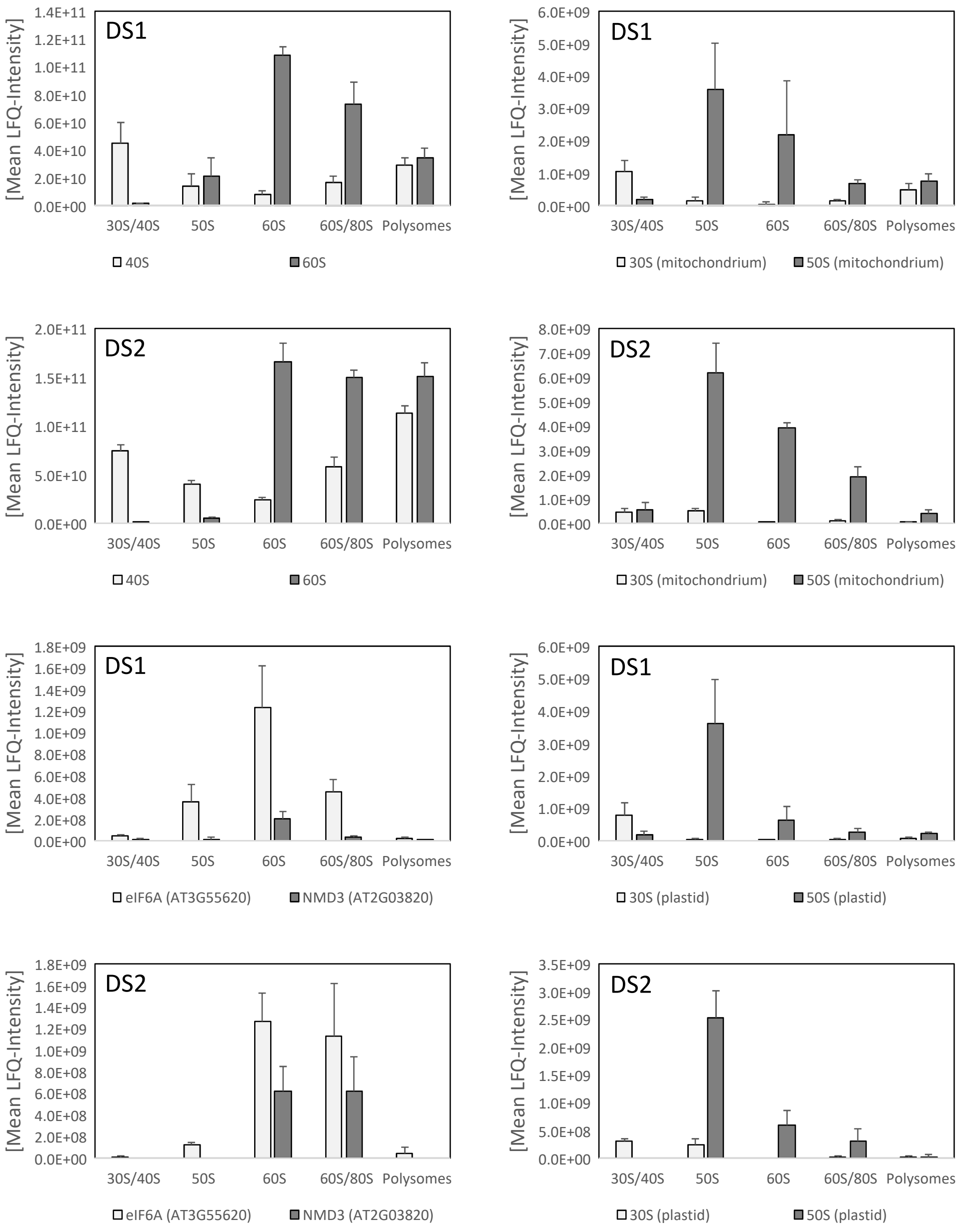

DS1
[Mean LFQ-Intensity]
1.4E+11
1.2E+11
1.0E+11
8.0E+10
6.0E+10
4.0E+10
2.0E+10
0.0E+00
30S/40S
50S
60S
60S/80S
Polysomes
40S
60S
DS1
[Mean LFQ-Intensity]
6.0E+09
5.0E+09
4.0E+09
3.0E+09
2.0E+09
1.0E+09
0.0E+00
30S/40S
50S
60S
60S/80S
Polysomes
30S (mitochondrium)
50S (mitochondrium)
DS2
[Mean LFQ-Intensity]
2.0E+11
1.5E+11
1.0E+11
5.0E+10
0.0E+00
30S/40S
50S
60S
60S/80S
Polysomes
40S
60S
DS2
[Mean LFQ-Intensity]
8.0E+09
7.0E+09
6.0E+09
5.0E+09
4.0E+09
3.0E+09
2.0E+09
1.0E+09
0.0E+00
30S/40S
50S
60S
60S/80S
Polysomes
30S (mitochondrium)
50S (mitochondrium)
DS1
[Mean LFQ-Intensity]
1.8E+09
1.6E+09
1.4E+09
1.2E+09
1.0E+09
8.0E+08
6.0E+08
4.0E+08
2.0E+08
0.0E+00
30S/40S
50S
60S
60S/80S
Polysomes
eIF6A (AT3G55620)
NMD3 (AT2G03820)
DS1
[Mean LFQ-Intensity]
6.0E+09
5.0E+09
4.0E+09
3.0E+09
2.0E+09
1.0E+09
0.0E+00
30S/40S
50S
60S
60S/80S
Polysomes
30S (plastid)
50S (plastid)
DS2
[Mean LFQ-Intensity]
1.8E+09
1.6E+09
1.4E+09
1.2E+09
1.0E+09
8.0E+08
6.0E+08
4.0E+08
2.0E+08
0.0E+00
30S/40S
50S
60S
60S/80S
Polysomes
eIF6A (AT3G55620)
NMD3 (AT2G03820)
DS2
[Mean LFQ-Intensity]
3.5E+09
3.0E+09
2.5E+09
2.0E+09
1.5E+09
1.0E+09
5.0E+08
0.0E+00
30S/40S
50S
60S
60S/80S
Polysomes
30S (plastid)
50S (plastid)

**Supplemental Figure S7.** Proteomic characterization of ribosome preparations obtained from root material by sucrose density gradient fractionation experiments DS1 and DS2, i.e. the root subset of Pride repository data set PXD016292. Ribosome complexes are represented by sums of LFQ-intensities of all detected RPs of 40S, 60S, and mitochondrial or plastid 30S and 50S subunits within each fraction. Non-translating immature 60S subunits are assessed by Arabidopsis homologs eIF6A and NMD3 of yeast cytosolic 60S maturation factors, means +/- standard error of the four fractionations of DS 1 or DS2 (Supplemental Table S4).

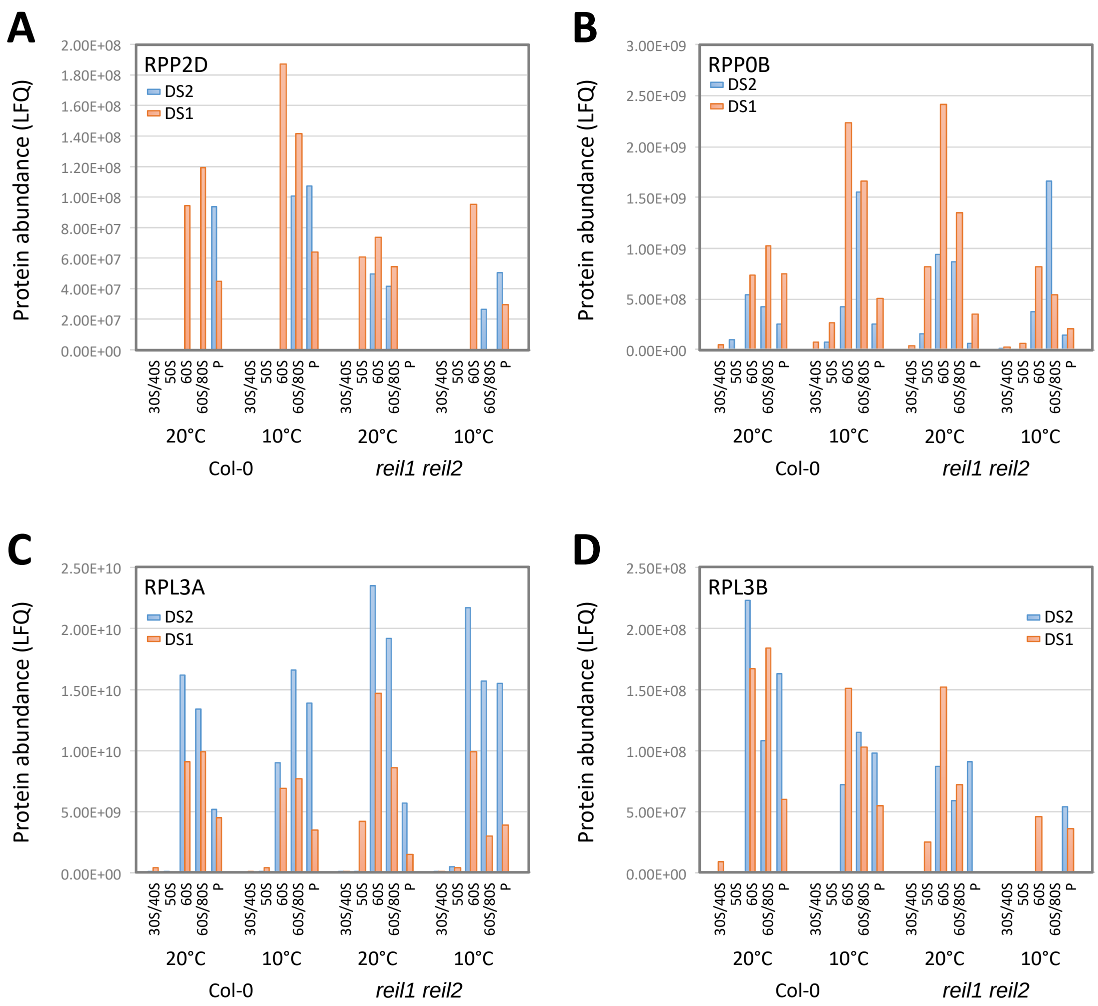

A
RPP2D
DS2
DS1
Protein abundance (LFQ)
2.00E+08
1.80E+08
1.60E+08
1.40E+08
1.20E+08
1.00E+08
8.00E+07
6.00E+07
4.00E+07
2.00E+07
0.00E+00
B
RPP0B
3.00E+09
2.50E+09
2.00E+09
1.50E+09
1.00E+09
5.00E+08
0.00E+00
C
RPL3A
2.50E+10
2.00E+10
1.50E+10
1.00E+10
5.00E+09
0.00E+00
D
RPL3B
2.50E+08
2.00E+08
1.50E+08
1.00E+08
5.00E+07
0.00E+00
30S/40S
50S
60S
60S/80S
P
20°C
10°C
Col-0
reil1 reil2

**Supplemental Figure S8.** Distribution analysis of RPP2D **(A)**, RPP0B **(B)**, RPL3A **(C)**, and RPL3B **(D)** across the sampled ribosome fractions and analyzed conditions, DS1 orange, DS2 blue.
